# Supervised Adversarial Alignment of Single-Cell RNA-seq Data

**DOI:** 10.1101/2020.01.06.896621

**Authors:** Songwei Ge, Haohan Wang, Amir Alavi, Eric Xing, Ziv Bar-Joseph

## Abstract

Dimensionality reduction is an important first step in the analysis of single cell RNA-seq (scRNA-seq) data. In addition to enabling the visualization of the profiled cells, such representations are used by many downstream analyses methods ranging from pseudo-time reconstruction to clustering to alignment of scRNA-seq data from different experiments, platforms, and labs. Both supervised and unsupervised methods have been proposed to reduce the dimension of scRNA-seq. However, all methods to date are sensitive to batch effects. When batches correlate with cell types, as is often the case, their impact can lead to representations that are batch rather than cell type specific. To overcome this we developed a domain adversarial neural network model for learning a reduced dimension representation of scRNA-seq data. The adversarial model tries to simultaneously optimize two objectives. The first is the accuracy of cell type assignment and the second is the inability to distinguish the batch (domain). We tested the method by using the resulting representation to align several different datasets. As we show, by overcoming batch effects our method was able to correctly separate cell types, improving on several prior methods suggested for this task. Analysis of the top features used by the network indicates that by taking the batch impact into account, the reduced representation is much better able to focus on key genes for each cell type.

## 1 Introduction

Single-cell RNA sequencing (scRNA-seq) has revolutionized the study of gene expression programs [14, 26]. The ability to profile genes at the single-cell level has revealed novel specific interactions and pathways within cells [41], differences in the proportions of cell types between samples [16, 42], and the identity and characterization of new cell types [38]. Several biological tissues, systems, and processes have recently been studied using this technology [16, 42, 41].

While studies using scRNA-seq provide many insights, they also raise new computational challenges. One of the major challenges involves the ability to integrate and compare results from multiple scRNA-seq studies. There are several different commercial platforms for performing such experiments, each with their own biases. Furthermore, similar to other high throughput genomic assays, scRNA-seq suffers from batch effects which can make cells profiled in one lab look very different from the same cells profiled at another lab [37, 36]. Moreover, other types of high throughput transcriptomics profiling, including microscopy-based techniques, are also generating single cell expression datasets [39, 7]. The goal of fully utilizing these spatial datasets motivates the development of methods that can combine them with scRNA-seq when studying specific biological tissues and processes.

A number of recent methods have attempted to address this challenge by developing methods for aligning scRNA-seq data from multiple studies of the same biological system. Many of these methods rely on identifying nearest neighbors between the different datasets and using them as anchors. Methods that use this approach include Mutual Nearest Neighbors (MNN) [11] and Seurat [35]. Others including scVI and scAlign first embed all datasets into a common lower dimensional space. scVI encodes the scRNA-seq data with a deep generative model conditioning on the batch identifiers [23] while scAlign regularizes the representation between two datasets by minimizing the random walk probability differences between the original and embedding spaces. While these methods were successful for some datasets, here we show that they are not always able to correctly match all cell types. A key problem with these methods is the fact that they are unsupervised and rely on the assumption that cell types profiled by the different studies overlap. While this works for some datasets, it may fail for studies in which cells do not fully overlap or for those containing rare cell types. Unsupervised methods tend to group rare types with the larger types making it hard to identify them in a joint space.

Recent machine learning work has focused on a related problem termed “domain generalization” or “domain adaptation”. Methods developed for these problems attempt to learn representations of diverse data that are invariant to technical confounders [24, 4]. These methods have been used for multiple applications such as machine translation for domain specific corpus [3] and face detection [27]. Several of the methods proposed for domain adaptation rely on the use of adversarial methods [4, 9, 20, 40], which has been proved effective to align latent distributions. In addition to the original task such as classification, these methods apply a domain classifier upon the learned representations. The encoder network is used for both improving accurate classification while at the same time reducing the impact of the domain (by “fooling” a domain classifier). This is achieved by learning encoder weights that simultaneously perform gradient *descent* on the label classification task and gradient *ascent* on the domain classification task.

Here we extend these approaches, coupling them with Siamese network learning [19] for overcoming batch effects in scRNA-seq analysis. We define a “domain” in this paper as a standalone dataset profiled at a single lab using a single platform. We define “label” as the cell type for each cell in the dataset. Considering the specificity of the cell types in the scRNA-seq datasets, we propose a conditional pair sampling strategy that constrains input pair selection when training the adversarial network. We discuss how to formulate a domain adaptation network for scRNA-seq data, how to learn the parameters for the network, and how to train it using available data.

We tested our method on several datasets ranging in size from 10 to 39 cell types and from 4 to 155 batches. As we show, for all of the datasets our domain adversarial method improves on previous methods, in some cases significantly. Visualization of the learned representation from several different methods helps highlight the advantages of the domain adversarial framework. As we show, the framework is able to accurately mitigate the batch effects while maintaining the grouping of cells from the same type across different batches. Biological analysis of the resulting model identifies key genes that can correctly distinguish between cell types across different experiments. Such batch invariant genes are promising candidates for a cell type specific signature that can be used across different studies to annotate cells.

## 2 Methods

### 2.1 Problem Formulation

To formulate the problem we start with a few notation definitions. We assume that the single cell RNA-seq data are drawn from the input space **X** ∈ R*^p^* where each sample (a cell) **x** has *p* features corresponding to the gene expression values. Cells are also associated with the label *y* ∈ **Y** = 1, 2*, …, K* which represents their cell types. We associate each sample with a specific domain/batch *d* ∈ 𝒟 that represents any standalone dataset profiled at a single lab using a single platform. Note that we will use domain and batch interchangeably in this paper for convenience. The data are divided into a training set and a test set that are drawn from multiple studies. The domains used to collect training data are not used for the test set and so batch effects can vary between the training and test data. In practice, each of the domains only contains a small subset of the cell types. This means that the distribution of cell types is correlated with the distribution of domains. Thus, the methods that naively learn cell types based on expression profile [2, 17, 21] may instead fit domain information and not generalize well to the unobserved studies.

### 2.2 Domain Adversarial Training with Siamese Network

To overcome this problem and remove the domain impact when learning a cell type representation we propose a neural network framework which includes three modules as shown in Figure 1: scRNA encoder, label classifier, and domain discriminator. The encoder module *f_e_*(**x**; *θ_e_*) is used to reduce the dimensions of the data and contains fully connected layers which produce the hidden features, where *θ_e_* represents the parameters in these layers. The label classifier *f_l_*(*f_e_*; *θ_l_*) attempts to predict the label of input **x_1_** whereas the goal of the domain discriminator *f_d_*(*f_e_*; *θ_d_*) is to determine whether a pair of inputs **x_1_** and **x_2_** are from the same domain or not. Past work for classifying scRNA-seq data only attempted to minimize the loss function for the label classifier 𝓛*_l_*(*f_l_*(*f_e_*; *θ_l_*)) [22, 2]. Here, we extend these methods by adding a regularization term based on the adversarial loss of the domain discriminator *L_d_*(*f_d_*(*f_e_*; *θ_d_*)) which we will elaborate later.

The overall loss *E* on a pair of samples **x_1_** and **x_2_** is denoted by:

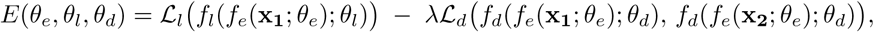

where *λ* can control the trade-off between the goals of domain invariance and higher classification accuracy. For convenience, we use **z_1_** and **z_2_** to denote the hidden representations of **x_1_** and **x_2_** calculated from *f_e_*(**x**; *θ_e_*). Inspired by Siamese networks [19], we implement our domain discriminator by adopting a contrastive loss [10]:

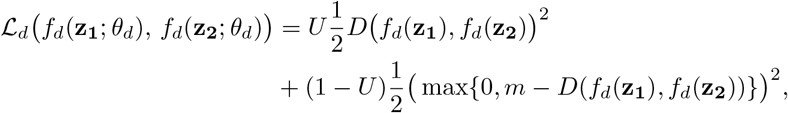

where *U* = 1 indicates that two samples are from the same domain *d* and *U* = 0 indicates that they are not, *D*(·) is the euclidean distance, and *m* is the margin that indicates the prediction boundary. The domain discriminator parameters, *θ_d_*, are updated using back propagation to *maximize* the total loss *E* as mentioned in the Introduction while the encoder and classifier parameters, *θ_e_* and *θ_l_*, are updated to *minimize E*. To allow all three modules to be updated together end-to-end, we use a Gradient Reversal Layer (Figure 1) [9, 28]. Specifically, Gradient Reversal Layers (GRL) have no effect in forward propagation, but flip the sign of the gradients that flow through them during backpropagation. The following provides the overall optimization problems solved for the network parameters:

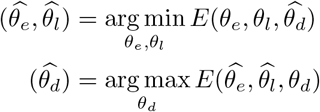

In other words, the goal of the domain discriminator is to tell if two samples are drawn from the same or different batches. By optimizing the scRNA encoder adversarially against the domain discriminator, we attempt to make sure that the network representation cannot be used to classify based on domain knowledge. During the training, the maximization and minimization tasks compete with each other, which is achieved by adjusting the representations to improve the accuracy of the label classifier and simultaneously fool the domain discriminator.

**Fig. 1:**
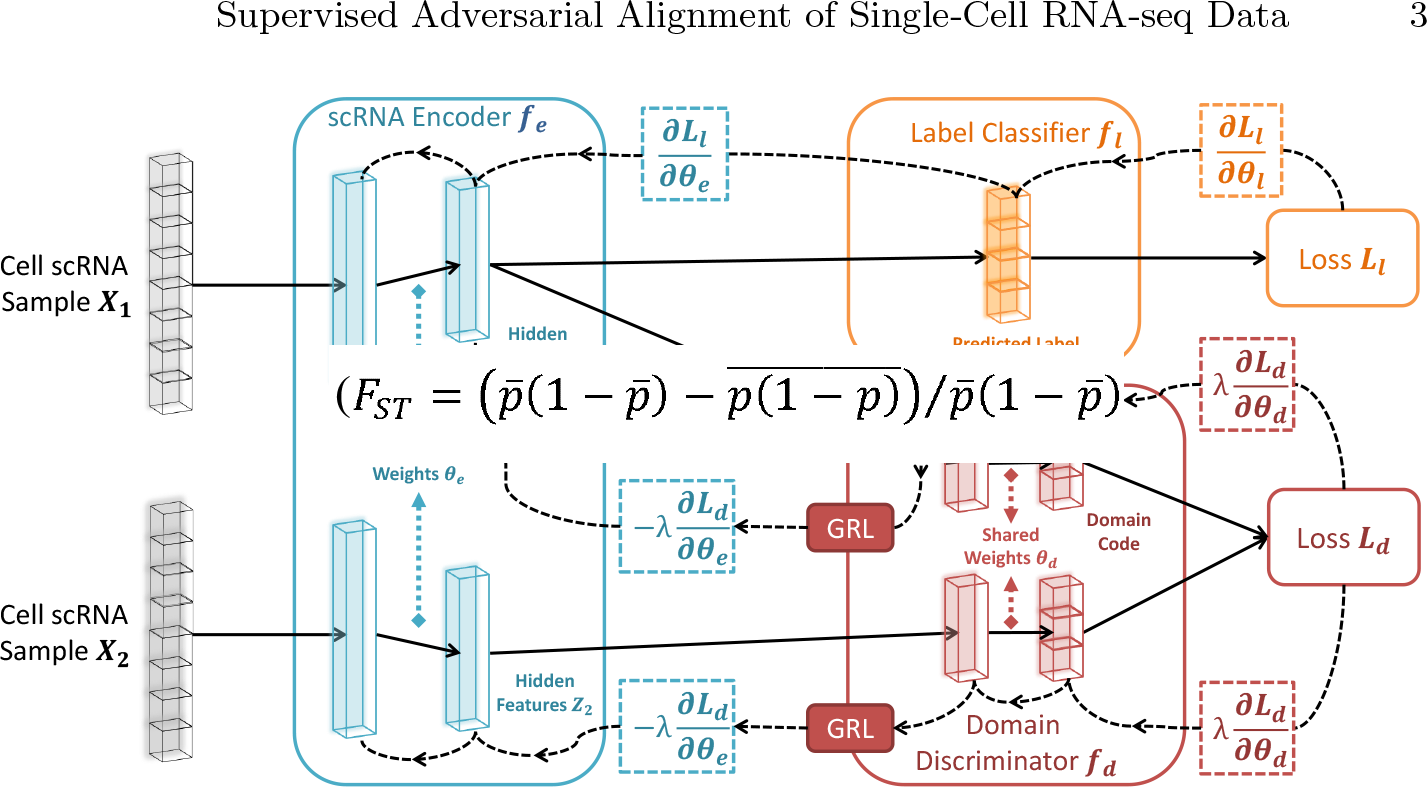
Architecture of scDGN. The network includes three modules: scRNA encoder *f_e_* (blue), label classifier *f_l_* (orange) and domain discriminator *f_d_* (red). Note that the red and orange networks use the same encoding as input. Solid lines represent the forward direction of the neural network while the dashed lines represent the backpropagation direction with the corresponding gradient it passes. Gradient Reversal Layers (GRL) have no effect in forward propagation, but flip the sign of the gradients that flow through them during backpropagation. This allows the combined network to simultaneously optimize label classification and attempt to “fool” the domain discriminator. Thus, the encoder leads to representations that are invariant to the different domains while still distinguishing cell types.

### 2.3 Conditional Domain Generalization Strategy

Most prior domain adaption or generalization methods focused on the cases where the distribution of labels is independent of the domains [24, 4]. In contrast, as we show in Results, for scRNA-seq experiments different studies tend to focus on certain cell types [16, 41, 42]. Consequently, it is not reasonable to completely merge the scRNA-seq data from different batches. To be specific, aligning the scRNA-seq data from two batches with different sets of cell types would sacrifice its biological significance and prevent the cell classifier from predicting effectively. To overcome this issue, instead of arbitrarily choosing positive pairs (samples from the same domain) and negative pairs (samples from different domains), we constrain the selection as follows: 1) for positive pairs, only the samples with different labels from the same domain are selected. 2) for negative pairs, only the samples with the same label from different domains are selected. Figure 2 provides a visual interpretation of this strategy. Formally, letting *y_i_* and *z_i_* represent the *i*-th sample’s cell-type label and domain label respectively, we have the following equations to define the value of *U* for sample pairs:

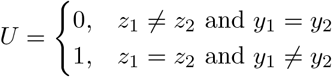

**Fig. 2:**
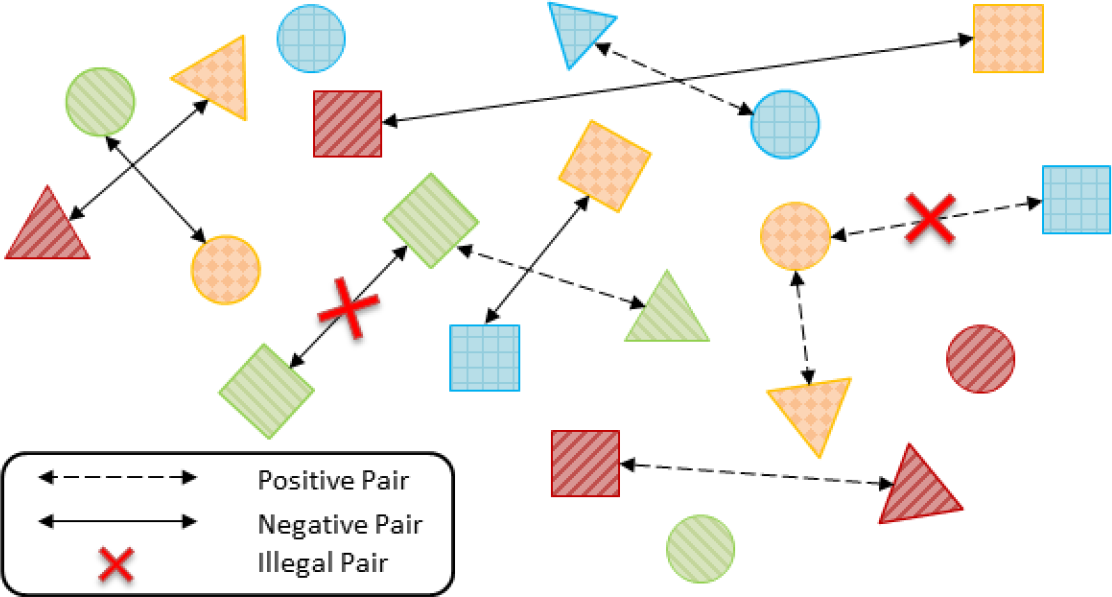
Conditional domain generalization strategy: Shapes represent different labels and colors (or patterns) represent different domains. For negative pairs from different domains, we only select those samples with the same label. For positive pairs from the same domain, we only select the samples with different labels.

This strategy prevents the domain adversarial training from aligning samples with different labels or separating samples with same labels. For example, in order to fool the discriminator with a positive pair, the encoder must implicitly increase the distance of two samples with different cell types. Therefore, combining this strategy with domain adversarial training allows the network to learn cell type specific, focused representations. We term our model Single Cell Domain Generalization Network (scDGN).

## 3 Results

### 3.1 Experiment Setups

#### Datasets

To test our method and to compare it to previous methods for aligning and classifying scRNA-seq data, we used several recent datasets. These datasets contain between 6,000 and 45,000 cells, and all include cells profiled in multiple experiments by different labs and on different platforms.

#### scQuery

We use a subset of the dataset provided by scQuery, which includes uniformly processed data from hundreds of studies [2]^1^. The dataset we use contains 44,490 samples from 155 different experiments. scQuery assigns cells to 39 types spanning diverse categories ranging from immune cells to neurons to organ specific cells. We use 99 of the 155 batches for training, 26 for validation, and 30 for testing. We provide a list of the studies used for each set in the Appendix A.1. Statistics for the different datasets are shown in Table 1. RPKM normalization is applied to the 20,499 genes in each sample. Note that while there are 39 cell-types in the training set, only 19 and 23 of them are included in the validation and test set. This mimics the application of the methods to future studies that may not profile all types of cells.

**Table 1:** Basic statistics for scQuery, Suerat pancreas, and PBMC datasets

|  | scQuery |  |  | Seurat pancreas |  |  | Seurat pbmc |  |  |
| --- | --- | --- | --- | --- | --- | --- | --- | --- | --- |
|  | data | cell type | domain | data | cell type | domain | data | cell type | domain |
| training | 37697 | 39 | 99 | 6321 | 13 | 3 | 25977 | 10 | 8 |
| validation | 3023 | 19 | 26 | - | - | - | - | - | - |
| test | 3770 | 23 | 30 | 638 | 13 | 1 | 2992 | 10 | 1 |

#### PBMC

The Peripheral Blood Mononuclear Cells (PBMC) dataset contains 28,969 cells assigned to 10 blood cell types. The data are profiled in 9 batches (from 8 different sequencing technologies) [5]. We use the data from the *10xChromiumv2A* platform as test data and the rest as training data. Following the provided tutorial [1], we use the top 3000 variable genes for the analysis.

#### Seurat pancreas

The Seurat pancreas dataset is designed for evaluating single cell alignment algorithms and contains 6321 scRNA-seq samples of human pancreatic islet cell produced by four studies. We use the smallest study for the test data and the other three for training as shown in Table 1. Thirteen canonical labels of the pancreatic islet cell are assigned to cells in each study. Similar to the Seurat PBMC dataset, we only used the 3000 most variable genes. To further simulate the correlation between cell types and domains for this dataset we randomly remove the data for 6 of the 13 cell types for each of the training domains. As a result, we construct 6 synthetic datasets based this strategy to evaluate the alignment performance of different methods under a high label-domain correlation setting. The specific cell type information of each dataset is listed Appendix A.3.

#### Model Configurations

We used the network of Lin et al [22] as the components for the encoder and the label classifier in our model. The encoder contains two hidden layers with 1136 and 100 units. The label classifier is directly connected to the 100 unit layer and makes predictions based on these values. The domain discriminator contains an additional hidden layer with 64 units and is also connected to the 100 unit layer of the encoder (Figure 1). For each layer, *tanh*() is used as the non-linear activation function. We test several other possible configurations but did not observe improvement in performance. As is commonly done, we use a validation set to tune the hyperparameters for learning including learning rates, decay, momentum, and the adversarial weight and margin parameters *λ* and *m*. Generally, our analysis indicates that for larger datasets a lower weight *λ* and larger margin *m* for the adversarial training is preferred and vice versa. More details about the hyperparameters and training are provided in Appendix A.2.

#### Baselines

We compared scDGN to several prior methods for classifying and aligning scRNA-seq data. These included the neural network (NN) model of Lin et al [22] which is developed for classifying scRNA-seq data, CaSTLe [21] which performs cell type classification based on transfer learning, and several state-of-the-art alignment methods. For alignment, we compared to MNN [11] which utilizes mutual nearest neighbors to align data from different batches, scVI [23] which trains a deep generative model on the scRNA-seq data and uses an explicit batch identifier to retain conditional independence property of the representation, and Seurat [35] which first identifies the anchors among different batches and then projects different datasets using a correction vector based on the order defined by hierarchical clustering with pairwise distances. Our comparisons include both visual projection of the learned alignment (Figure 4 and 5) and quantitative analysis of the accuracy of the predicted test cell types (Table 2). For the latter, to enable comparisons of the supervised and unsupervised methods, we used the resulting aligned data from the unsupervised methods to train a neural network that has the same configuration as Lin et al [22]. For scVI, which results in a much lower dimensional representation, we used a smaller input vector and a smaller hidden layer. Note that these alignment methods actually use the scRNA-seq test data to determine the final dimensionality reduction function while our method does not utilize the test data for any model decision or parameter learning. To effectively apply Seurat to scQuery, we remove the batches which have *<* 100 samples. Also, for those datasets that the assumption of overlapped cell types is not guaranteed such as scQuery, we find that the performance of MNN highly depends on the order of alignment. Therefore, for MNN on the scQuery dataset, we use 10 random permutations of batch orders and report the average accuracy.

**Fig. 4:**
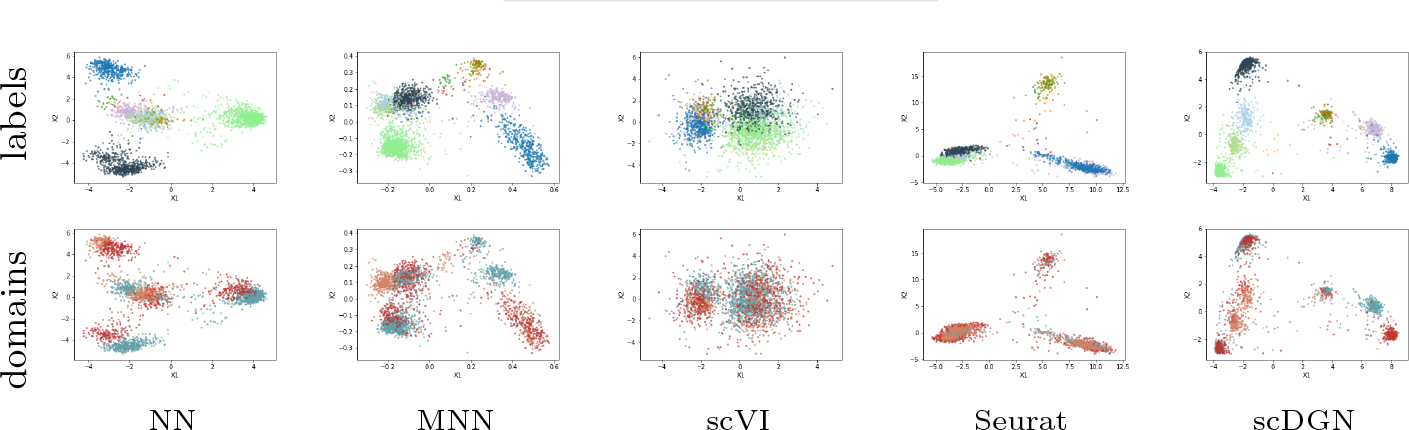
PCA visualizations of the representations learned by different models on the full Pancreas2 dataset. Colors for different cell types and domains are shown in the legend at the top.

**Table 2:** Overall performances of different methods. *MI* represents the mutual information between batch and cell type in the corresponding dataset. The highest test accuracy for each dataset is bolded.

| Experiments | MI | NN[22] | CaSTLe [21] | MNN [11] | scVI [23] | Seurat [35] | scDGN |
| --- | --- | --- | --- | --- | --- | --- | --- |
| scQuery | 3.025 | 0.255 | 0.156 | 0.200 | 0.257 | 0.144 | **0.286** |
| PBMC | 0.112 | 0.861 | 0.865 | 0.859 | 0.808 | 0.830 | **0.868** |
| Pancreas 1 | 0.902 | 0.720 | 0.705 | 0.591 | 0.855 | 0.812 | **0.856** |
| Pancreas 2 | 0.733 | 0.891 | 0.764 | 0.764 | 0.852 | 0.825 | **0.918** |
| Pancreas 3 | 0.931 | 0.545 | 0.722 | 0.722 | 0.651 | **0.751** | 0.663 |
| Pancreas 4 | 0.458 | 0.927 | 0.914 | 0.914 | **0.925** | 0.881 | **0.925** |
| Pancreas 5 | 0.849 | 0.928 | 0.882 | **0.932** | 0.895 | 0.865 | 0.923 |
| Pancreas 6 | 0.670 | 0.944 | 0.917 | 0.946 | 0.893 | 0.907 | **0.950** |
| Average | - | 0.826 | 0.817 | 0.842 | 0.845 | 0.840 | **0.872** |

**Table 4:** GO analysis results for top 100 scQuery liver genes in the scDGN method.

| term_name | term_id | $p_{adj}$ | $-\log_{10} p_{adj}$ |
| --- | --- | --- | --- |
| chylomicron remodeling | GO:0034371 | 3.04042E-05 | 4.517066786 |
| positive reg. of cholesterol esterification | GO:0010873 | 3.04042E-05 | 4.517066786 |
| negative reg. of cellular component organization | GO:0051129 | 3.94437E-05 | 4.404022507 |
| protein-lipid complex remodeling | GO:0034368 | 7.34551E-05 | 4.133978335 |
| plasma lipoprotein particle remodeling | GO:0034369 | 7.34551E-05 | 4.133978335 |
| protein-containing complex remodeling | GO:0034367 | 8.8522E-05 | 4.052948555 |

### 3.2 Overall Performance

As mentioned above, we use the validation set to select the best model when using the scQuery dataset. For the smaller datasets, we use the model obtained after 250 epochs (all models converged after this number of epochs). Test accuracy for the different methods is presented in Table 2. We show both mean and standard deviation of the accuracy for 10 randomly initialized experiments. We also report the performances on different cell types in Appendix B. In addition, Table 2 presents the Mutual Information (MI) between labels and domains which corresponds to the difficulty of the dataset. A larger MI indicates that models that do not account for the domain are likely to fit the domain information rather than the cell type. For the scQuery dataset, we find the accuracy is low for all methods indicating that this dataset is relatively difficult. This is corroborated by the large MI value. For such data we see a clear advantage for the scDGN: scDGN improves by over 10% over all other methods (*p* = 5.069 10*^−^*^5^ based on Student’s t-test when compared to the NN baseline which is tied for second best). The improvements over other single cell alignment methods are even more significant. scDGN also achieves the best performance on the second largest dataset, the PBMC dataset. However, given the very low MI for this dataset the performance of the other methods, including the baseline NN, is almost as good as the performance of scDGN. The third dataset we test on is the Seurat pancreas dataset. This is the smallest dataset and so it has the least number of training samples. Still, of the 6 settings we tested (which differed in the subset of cells that were excluded from training), we find that scDGN is the top performer in 4 of them, comparable to the top performer for another 1 and in only one setting (Pancreas 3, with the highest MI) is significantly outperformed by Seurat. Note that even for the Pancreas 3 data the domain adversarial training helps: using this the scDGN is able to improve by more than 20% over the baseline NN used for the label classifier.

### 3.3 Visualization of the Representation Learned by Alignment and Classification Methods

To further explore the effectiveness of the batch removal provided by our proposed domain adversarial training with conditional domain generalization strategy, we visualize the 100-dimensional hidden representations learned by NN and scDGN: Figure 3 presents both PCA and t-SNE plots for several different cell types across the three datasets. Points are colored using their batch IDs in order to evaluate batch effects. As can be seen, using scDGN we obtain results that are much better at mixing cells from the different batches when compared to the baseline NN model. The impact is larger for the pancreas datasets which have larger MI compared to the PBMC dataset, which helps explain the large increase in performance for these two datasets.

**Fig. 3:**
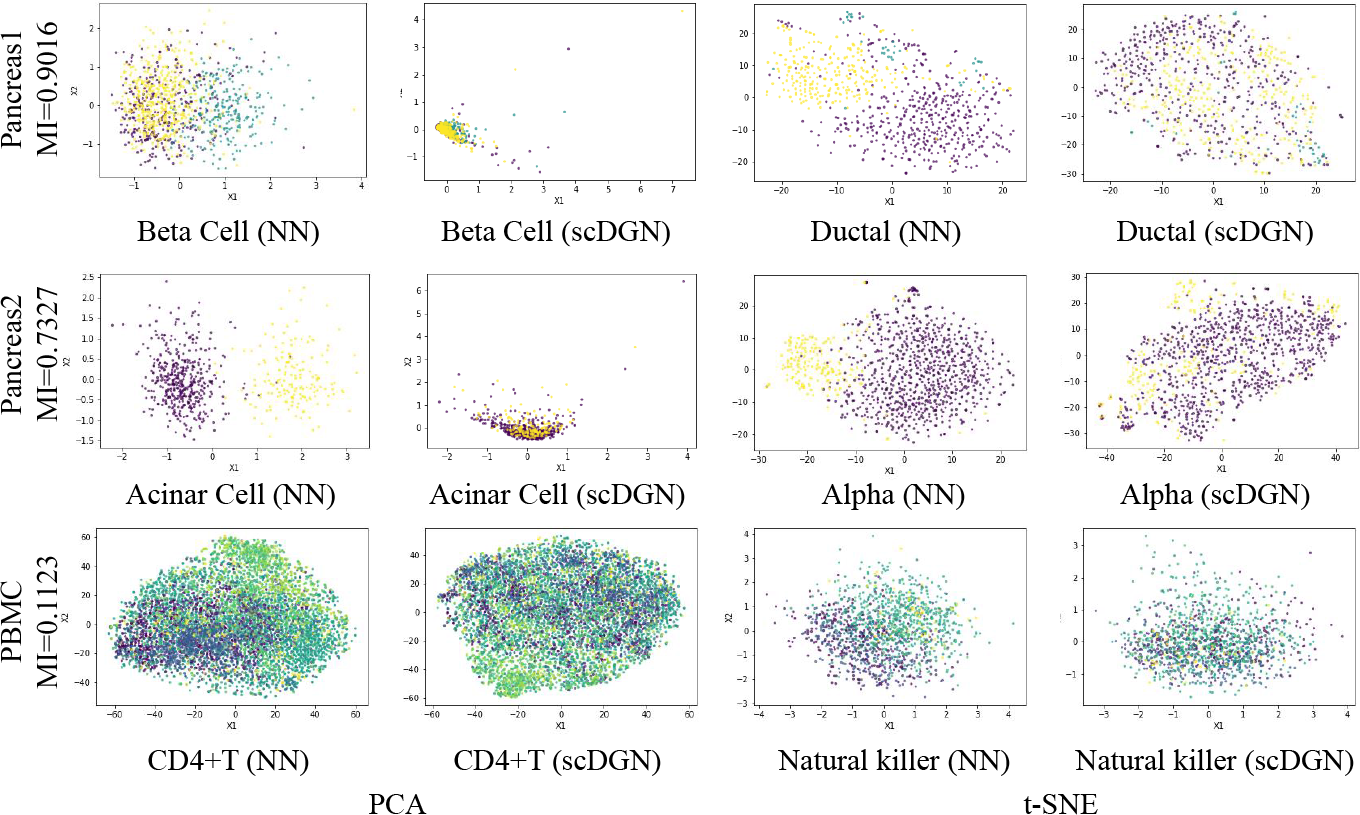
Visualization of learned representations for NN and scDGN: using PCA and t-SNE Rows: The three datasets we tested the method on. Columns: Methods and cell types. For each row, data from different batches are distinguished using different colors.

We next extended this comparison and visualized the learned (aligned) representations for all methods using data from both the Pancreas2 and scQuery datasets (Figures 4 and 5). For the Pancreas2 dataset, we visualize the entire dataset. For scQuery, given the large number of cell types and domains, we present PCA visualization of a subset of cell types and domains. As can be seen, in addition to scDGN, Seurat is also able to successfully mix the data from different batches. However, as the results in Table 2 indicate this may come at the expense of not correctly separating cell types. MNN and scVI are not always effective at removing batch effects for the cell types. In contrast, scDGN is able to do both domain mixing and cell type assignment, leading to its better performance overall. For example, for the acinar and alpha cell types in the pancreas dataset (Figure 4), only scDGN, MNN, and Seurat are able to align the data from different domains. However, MNN and Seurat over-correct the representation by aligning different cell types from different domains, mixing acinar and gamma cells. Additional visualizations for other cell types and domains can be found in Appendix C, where the same advantages of scDGN over other methods can be consistently observed.

**Fig. 5:**
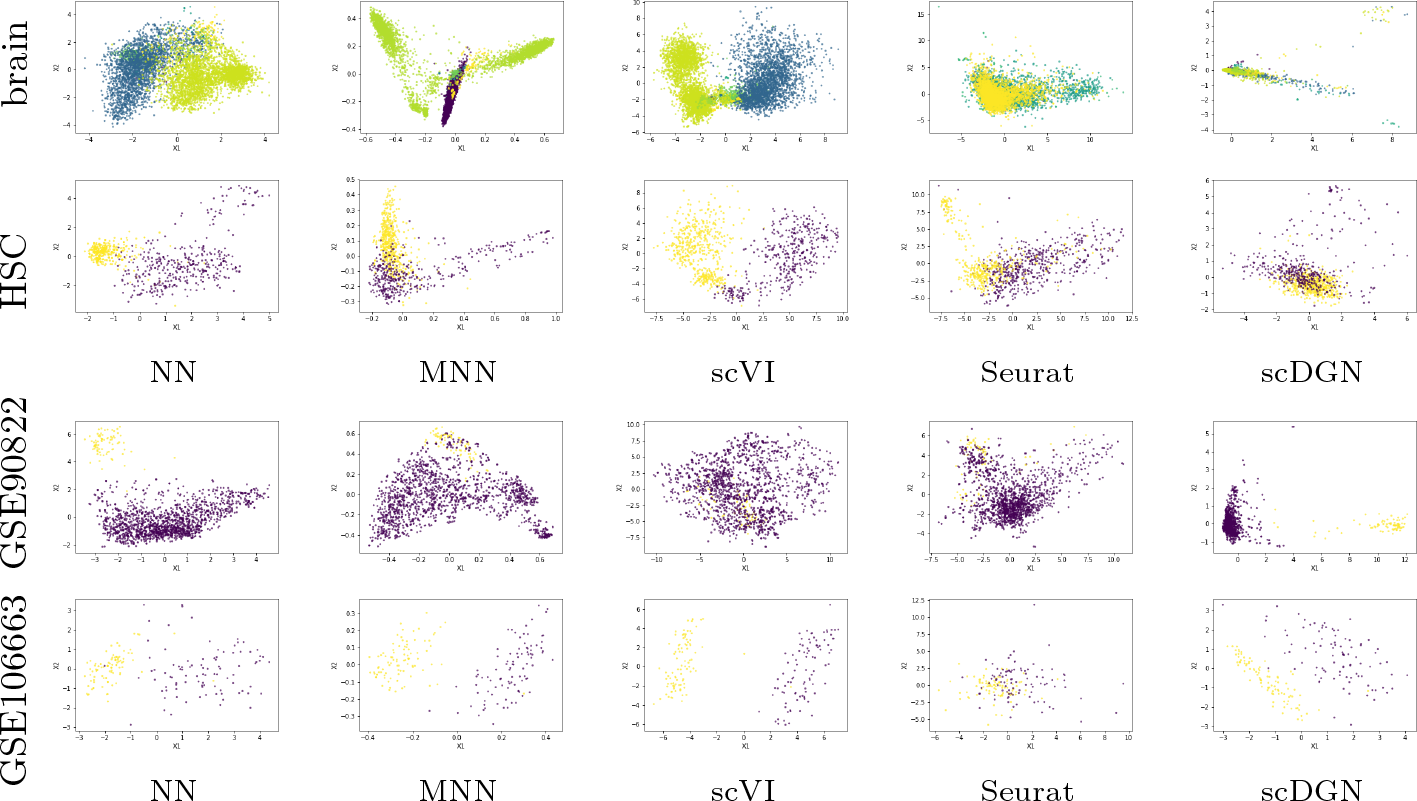
PCA visualizations of the representations of certain cell types and batches by different models for the scQuery dataset. Top two rows: Cell types. Colors represent different batches. HSC = hematopoietic stem cell. Bottom two rows: Batches. Colors represent different cell types.

### 3.4 Analysis of Key Genes

While NNs are often treated as black boxes, recent methods provide useful directions for making them more interpretable [30]. Here we use activation maximization, which relies on the gradient of the correct category logit with respect to the input vector to select the key inputs for each of the models [8, 32, 33]. Formally, given a particular cell type *i* and a trained neural network *ϕ*, activation maximization looks for important input genes *x^l^* by solving the following optimization problem:

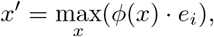

where *e_i_* is the natural basis vector associated with the *i*-th category. This can be solved through backpropagation, where the gradient of *ϕ*(*x*) with respect to *x*, which can be viewed as the weight of the first-order Taylor expansion of the neural network, are calculated to iteratively update the input. We follow a previous method [32] and initialize the optimization with a zero vector. Given this setting, we ran the optimization for 100 iterations with learning rate set to 1. The important genes are selected as those inputs leading to the largest changes compared with the initialization values. To compare scDGN and NN for certain cell types, we select the top *k* genes with the largest changes and perform GO analysis on these selected genes.

As an example, consider the genes identified for the liver cell type using the scQuery dataset. We select the top 100 genes for this cell type from NN and scDGN and present the enriched GO categories on Biological Process with adjusted p-value *<* 1.0 × 10*^−^*^4^ in Tables 3 and 4. We also list these genes by order in Appendix D.1. As can be seen, while a number of significant GO categories are identified for the top 100 NN genes, these are generic and not liver specific. They include general terms related to interactions between organs and immune response categories that are active in multiple organs and cell types. In sharp contrast, the categories identified for scDGN are much more specific and highlight key pathways that are mainly utilized in the liver. For example, the top category for the scDGN genes, “chylomicron remodeling”, refers to the main physiological purpose of chlyomicron remnants: to facilitate the return of bile lipoproteins and cholesterol to the liver [29]. Specifically, in this pathway chylomicrons (lipoproteins) are broken down (remodeled via hydrolysis) and converted to a form called “chlyomicron remnant” that is taken up by specific receptors that exist primarily on the surface of liver cells [12]. The second term, “pos. regulation of cholesterol esterification” refers to cholesterol esterification, a critical step in reverse cholesterol transport, the process in which excess cholesterol is sent to the liver to be removed from the body [15, 25]. Furthermore, Cholesteryl Ester Transfer Protein (CETP) is a key enzyme involved in this process and is highly expressed in liver cells, and variants of CETP are associated with increased risk of atherosclerosis [31, 15]. The fifth most significant term, “lipoprotein remodeling” is part of the two aforementioned processes. The top 100 genes identified by the scDGN include *apoa1* (main protein component of High-Density Lipoprotein cholesterol), *apoa2*, and *apoc1*, all of which encode lipoproteins that are primarily expressed in the liver [6, 18]. These genes were not included in the top 100 genes by the NN. We present the GO analysis results comparison for several additional cell types in Appendix D.

**Table 3:**
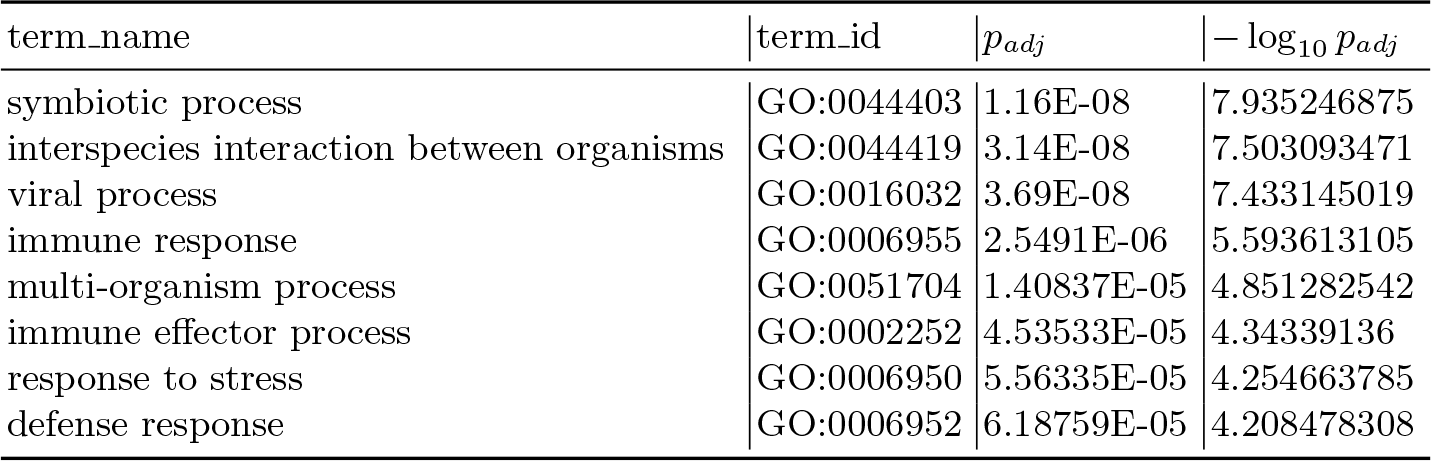
GO analysis results for top 100 scQuery liver genes in the NN method.

| term_name | term_id | $p_{adj}$ | $-\log_{10} p_{adj}$ |
| --- | --- | --- | --- |
| symbiotic process | GO:0044403 | 1.16E-08 | 7.935246875 |
| interspecies interaction between organisms | GO:0044419 | 3.14E-08 | 7.503093471 |
| viral process | GO:0016032 | 3.69E-08 | 7.433145019 |
| immune response | GO:0006955 | 2.5491E-06 | 5.593613105 |
| multi-organism process | GO:0051704 | 1.40837E-05 | 4.851282542 |
| immune effector process | GO:0002252 | 4.53533E-05 | 4.34339136 |
| response to stress | GO:0006950 | 5.56335E-05 | 4.254663785 |
| defense response | GO:0006952 | 6.18759E-05 | 4.208478308 |

## 4 Discussion

Single cell computational methods that do not account for batch effects are likely to fit the noise introduced by the batches. Several recent methods have been proposed for aligning scRNA-seq from multiple studies of the same tissues or processes. Most of these methods are unsupervised and assume that the cell types among different batches overlap. However, we show that these methods would fail on the studies in which cell types do not fully overlap, which is often the case when dealing with multiple datasets. To overcome this problem we extend a supervised scRNA-seq cell type assignment method based on NN and regularize its prediction to be invariant to batch effects.

Our method is based on the ideas of domain adversarial training. In such training, two competing tasks are used to optimize the representation of scRNA-seq data. The first focuses on the traditional goal of cell type identification while the second attempts to construct representations that are not affected by specific batch or experimental artifacts. This is accomplished by jointly minimizing a loss function that takes into account both goals, accounting for the weight of each of the goals using a gradient reversal layer. We also proposed a conditional strategy to avoid over-correction. We presented efficient learning methods for this setting and tested it on three large scale scRNA-seq datasets containing experiments from several different platforms for partially overlapping cell types.

As we show, our scDGN method is able to correctly identify cell types in the test (unobserved) datasets. For the largest dataset we tested on which contained close to 40 different cell types, scDGN significantly outperformed all prior methods. It also ranked first for the 2nd largest dataset and for all but 1 of the 6 tests on the third dataset. Importantly, it always outperformed the supervised learning based method indicating that batch effects should be addressed when designing such methods for cell type assignments. In addition to accurately assigning cell types, further analysis of significant genes indicates that by overcoming batch effects scDGN is better able to focus on relevant sets of genes for various cell types when compared to prior supervised methods, explaining its improvement in accuracy.

While scDGN performed best on the data we analyzed, there are a number of possible issues with this approach. First, it learns a large number of parameters which require large input datasets. However, as we showed, scDGN is able to perform well even for datasets with a few thousand cells which matches current sizes of scRNA-seq datasets. Second, scDGN is based on NNs which are often seen as a black box, making it hard to interpret the resulting model and its biological relevance. Recent work provides a number of directions that can be used to overcome this issue. As we showed, using activation maximization we were able to identify several relevant cell type specific genes in the learned network. Future work would include using additional NN interpretation methods, including LIME [30] or ROAR and KAR [13], to further identify the set of genes that play the largest role in the decisions the network makes. Third, as shown in Figure C.13, scDGN sometimes does not mix up the representations from different batches for all cell types. Considering the visualization results for NN in Figure C.18 and its competitive performance in Table 2 together, it may indicate that it is not always necessary to remove batch effects for the model to achieve high test accuracy. Therefore, it is worthwhile to further study when the alignment is imperative. Finally, unlike prior scRNA-seq alignment methods scDGN is supervised. While this is an advantage when it comes to accuracy, as we have shown, it may be a problem for the new data. We believe that as more scRNA-seq and other high throughput single cell data accumulate, we would have labeled data for most cell types which would enable training an scDGN for even more cell types. As we have shown with the scQuery dataset, for which scDGN significantly outperformed all other methods, when such data exists scDGN is able to correctly align experiments and platforms not seen in the training set.

scDGN is implemented in Python with the PyTorch API [34] and users can obtain the code and sampled data from https://github.com/SongweiGe/scDGN.

## A Experiment Details

### A.1 scQuery Dataset

The studies from which the data are collected for our scQuery dataset are shown as below:

**Training Set:** ERP017366, ERP021445, ERP022096, ERP022251, ERP022289, ERP022298, ERP022654, GSE101487, GSE101984, GSE102159, GSE102332, GSE102346, GSE102455, GSE102456, GSE103268, GSE103892, GSE104156, GSE104396, GSE105054, GSE106447, GSE106472, GSE106663, GSE107115, GSE107122, GSE108097, GSE109774, GSE109796, GSE112033, GSE115070, GSE22182, GSE29087, GSE33979, GSE38198, GSE42268, GSE42704, GSE42706, GSE45719, GSE52564, GSE57403, GSE57609, GSE59114, GSE60066, GSE61346, GSE61844, GSE62952, GSE64959, GSE64960, GSE65160, GSE66343, GSE66390, GSE66582, GSE68981, GSE69761, GSE69970, GSE70713, GSE71982, GSE72852, GSE72854, GSE72855, GSE72856, GSE75454, GSE75659, GSE75804, GSE76381, GSE77113, GSE78140, GSE78471, GSE78521, GSE79306, GSE79374, GSE79380, GSE79578, GSE79812, GSE80155, GSE80168, GSE80280, GSE80483, GSE81275, GSE84498, GSE87375, GSE89468, GSE90697, GSE90822, GSE90824, GSE90860, GSE92707, GSE93524, GSE94333, GSE94389, GSE94579, GSE98048, GSE98664, GSE98816, GSE98969, GSE98971, GSE99235, GSE99701, GSE99786, GSE99866.

**Validation Set:** ERP013319, ERP022703, GSE100120, GSE102163, GSE103267, GSE107053, GSE107527, GSE112642, GSE113043, GSE44183, GSE57393, GSE59127, GSE65924, GSE68770, GSE71585, GSE71794, GSE71802, GSE74534, GSE78401, GSE78510, GSE79108, GSE85234, GSE85627, GSE93421, GSE94388, GSE96981.

**Test Set:** ERP022293, GSE102827, GSE106471, GSE107740, GSE107909, GSE108291, GSE108478, GSE32190, GSE39522, GSE39523, GSE57249, GSE57391, GSE59129, GSE60749, GSE65525, GSE65970, GSE67120, GSE68769, GSE75790, GSE75901, GSE77705, GSE78045, GSE82174, GSE84324, GSE86479, GSE87631, GSE89900, GSE90047, GSE96986, GSE99058.

### A.2 Training Details

For the scQuery, we find that the optimal value of *λ* is 0.02 and *m* = 17. For other datasets we use *m* = 1 except for Pancreas 3 to which we apply *m* = 3. For Pancreas 3 we use *λ* = 0.1 and Pancreas 1 and 2 we use *λ* = 0.2. For all other datasets we use *λ* = 1. The models are implemented in PyTorch and trained on a machine with GeForce GTX 980 Ti and 32 GB memory. As for the ScQuery dataset, it takes around 9 and 19 seconds to train NN and scDGN for each epoch respectively.

### A.3 Seurat Pancreas Datasets

The cell type compositions in our pancreas datasets are shown as below:

|  | CelSeq | CelSeq2 | SMART-Seq2 | Fluidigm |
| --- | --- | --- | --- | --- |
| Pancreas 1 | gamma, ductal, mast, endothelial, beta, quiescent_stellate, macrophage | gamma, ductal, mast, epsilon, alpha, delta, acinar | macrophage, activated_stellate, schwann, epsilon, alpha, delta, acinar | All |
| Pancreas 2 | macrophage, activated_stellate, schwann, epsilon, alpha, delta, acinar | gamma, ductal, mast, endothelial, beta, quiescent_stellate, macrophage | gamma, ductal, mast, epsilon, alpha, delta, acinar | All |
| Pancreas 3 | gamma, ductal, mast, epsilon, alpha, delta, acinar | macrophage, activated_stellate, schwann, epsilon, alpha, delta, acinar | gamma, ductal, mast, endothelial, beta, quiescent_stellate, macrophage | All |
| Pancreas 4 | gamma, ductal, mast, epsilon, alpha, delta, acinar | gamma, ductal, mast, endothelial, beta, quiescent_stellate, macrophage | macrophage, activated_stellate, schwann, epsilon, alpha, delta, acinar | All |
| Pancreas 5 | macrophage, activated_stellate, schwann, epsilon, alpha, delta, acinar | gamma, ductal, mast, epsilon, alpha, delta, acinar | gamma, ductal, mast, endothelial, beta, quiescent_stellate, macrophage | All |
| Pancreas 6 | gamma, ductal, mast, endothelial, beta, quiescent_stellate, macrophage | macrophage, activated_stellate, schwann, epsilon, alpha, delta, acinar | gamma, ductal, mast, epsilon, alpha, delta, acinar | All |

## B Per Cell-type Performance

**Fig. B.1:**
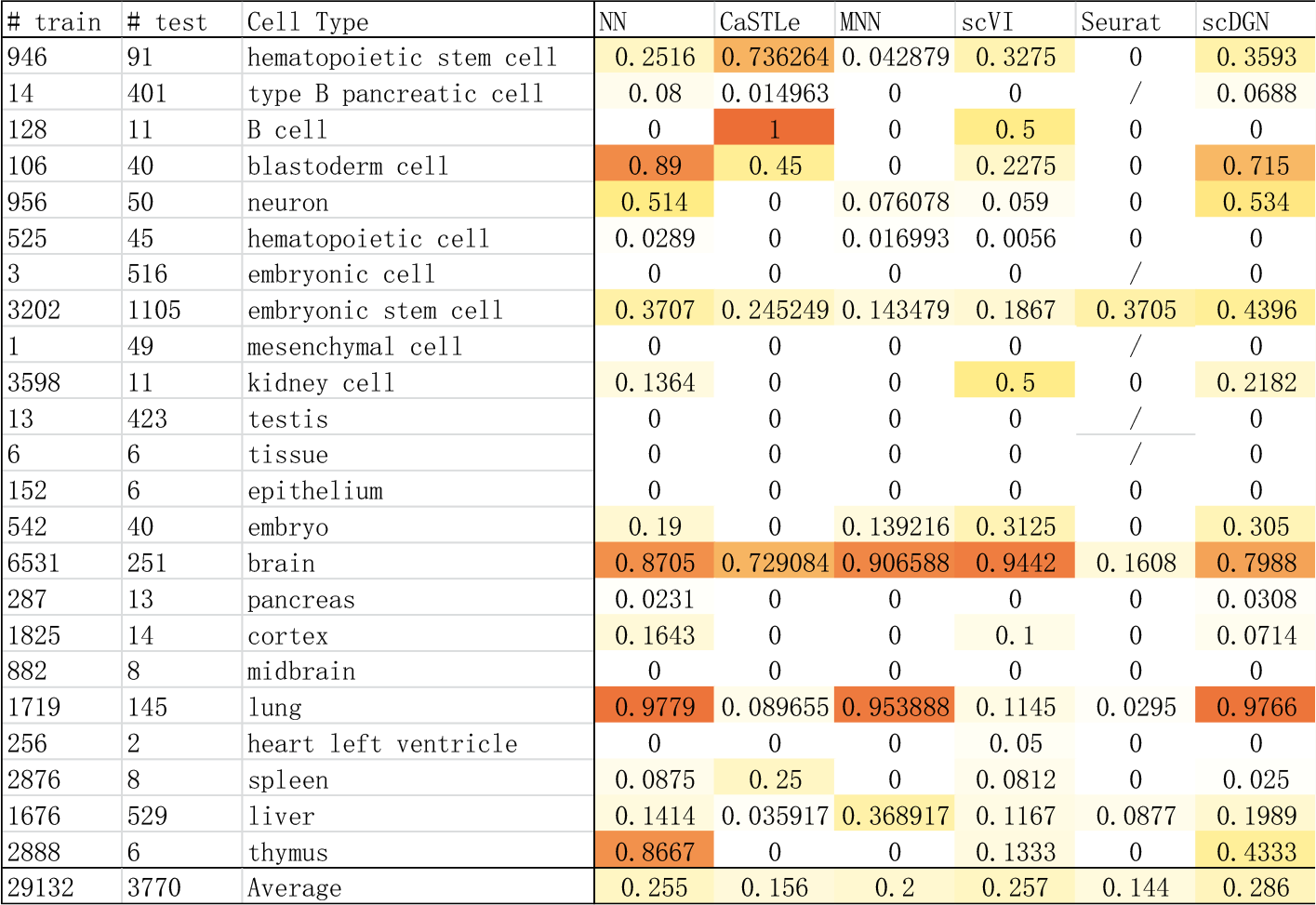
Test Accuracy of each model on different cell types from scquery dataset. The darker color represents the better performance. Note that the cell types that are not contained in the test dataset are not shown in this Table. The value is calculated by 10 experiments with different initializations for NN-based model to alleviate the randomness. The average is weighted by the number of the test samples. Note that scDGN performs the best on average.

**Fig. B.2:**
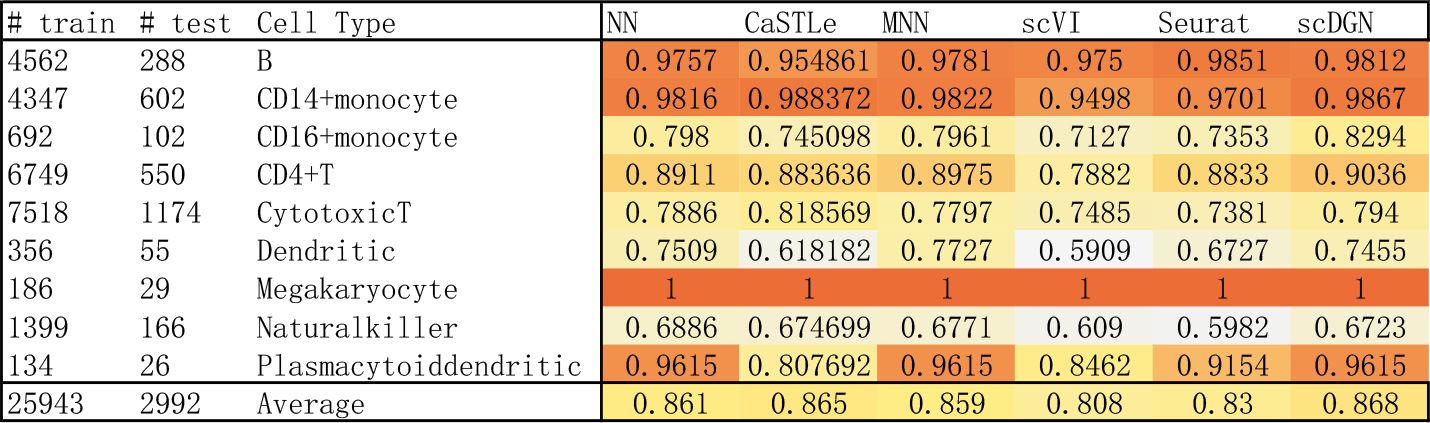
Test Accuracy of each model on different cell types from PBMC dataset.

**Fig. B.3:**
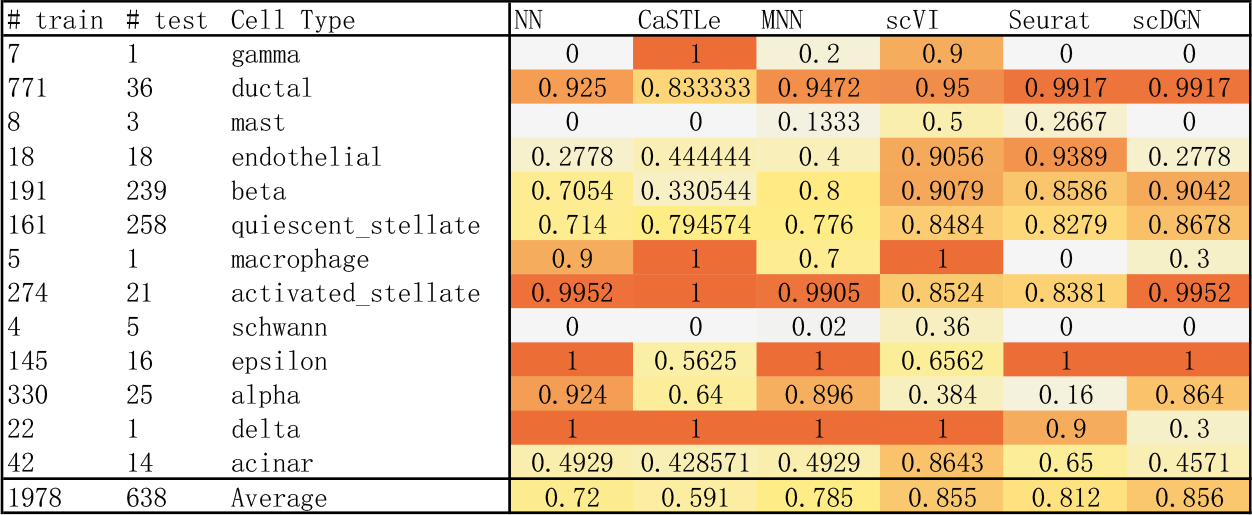
Test Accuracy of each model on different cell types from pancreas1 dataset.

**Fig. B.4:**
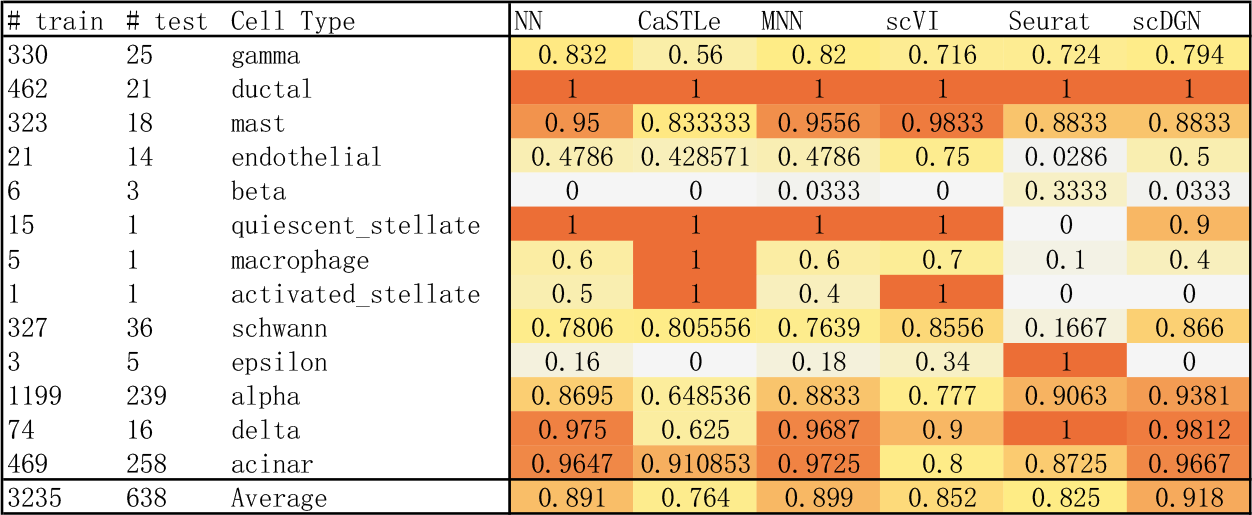
Test Accuracy of each model on different cell types from pancreas2 dataset.

**Fig. B.5:**
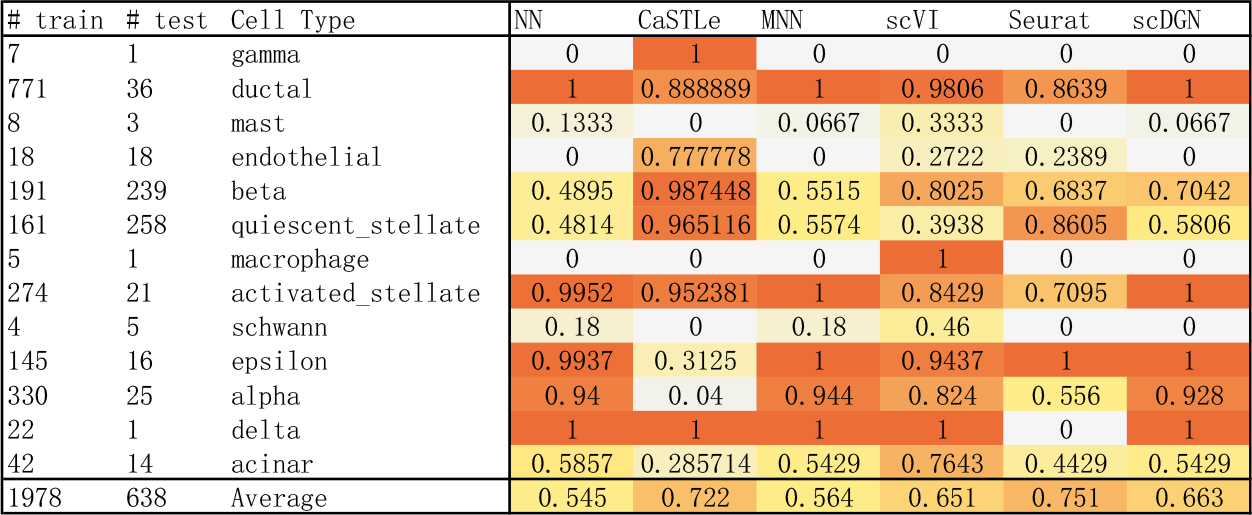
Test Accuracy of each model on different cell types from pancreas3 dataset.

**Fig. B.6:**
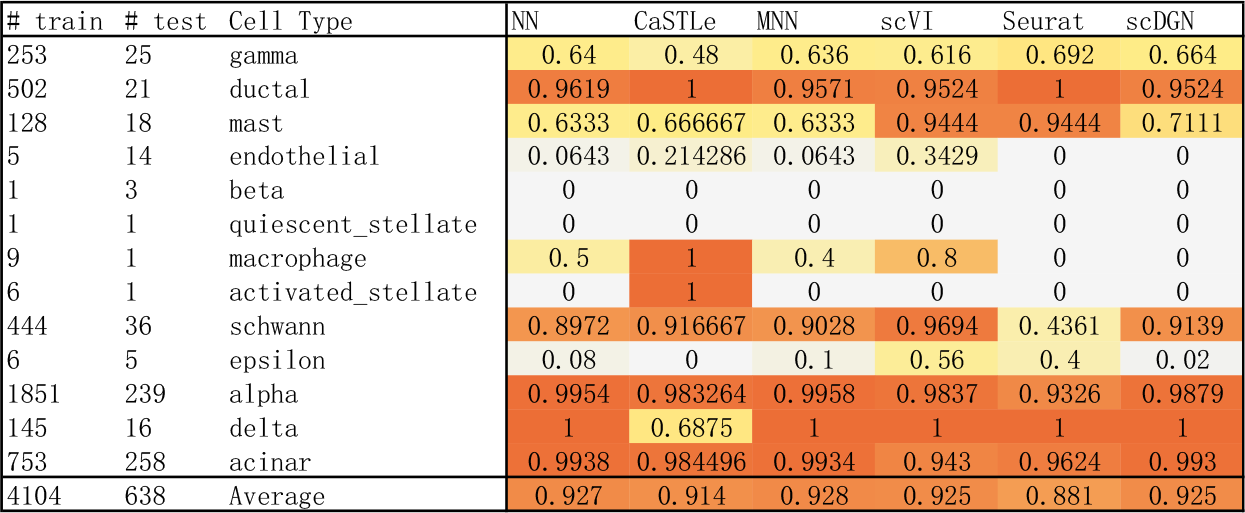
Test Accuracy of each model on different cell types from pancreas4 dataset.

**Fig. B.7:**
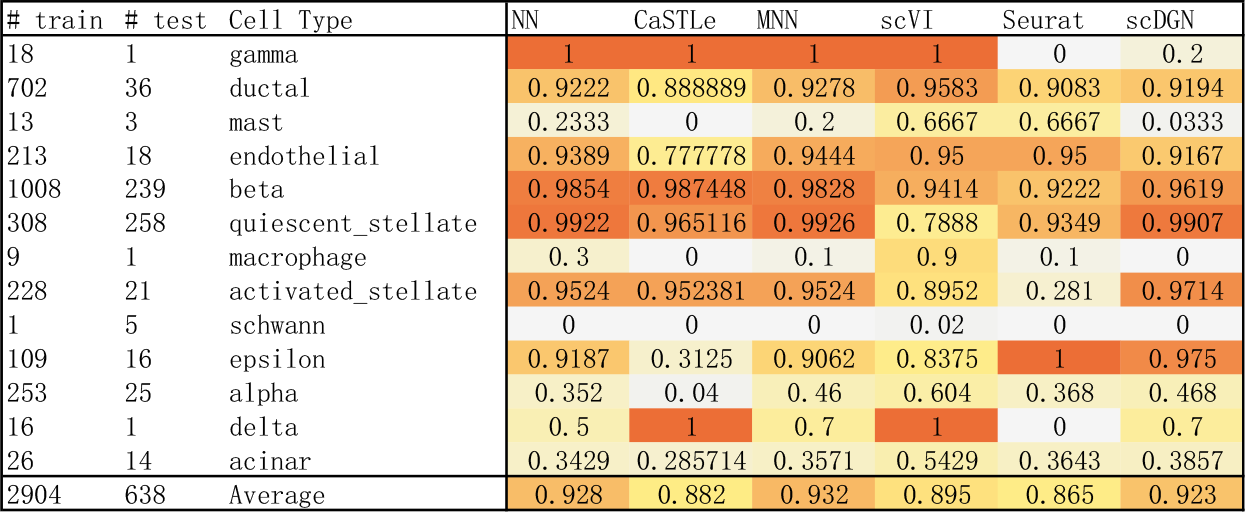
Test Accuracy of each model on different cell types from pancreas5 dataset.

**Fig. B.8:**
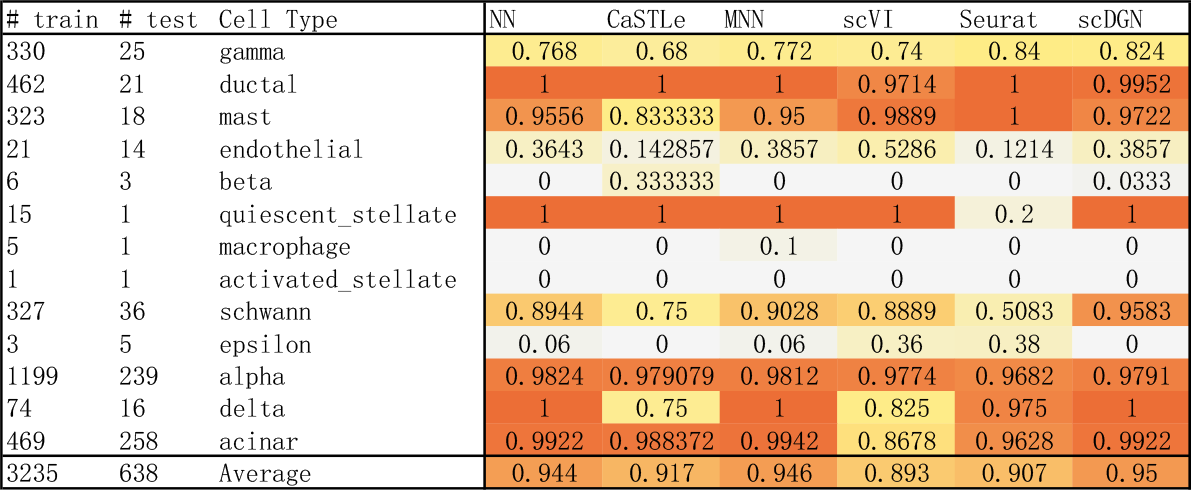
Test Accuracy of each model on different cell types from pancreas6 dataset.

## C Representation Visualization

### C.1 scQuery

**Fig. C.9:**
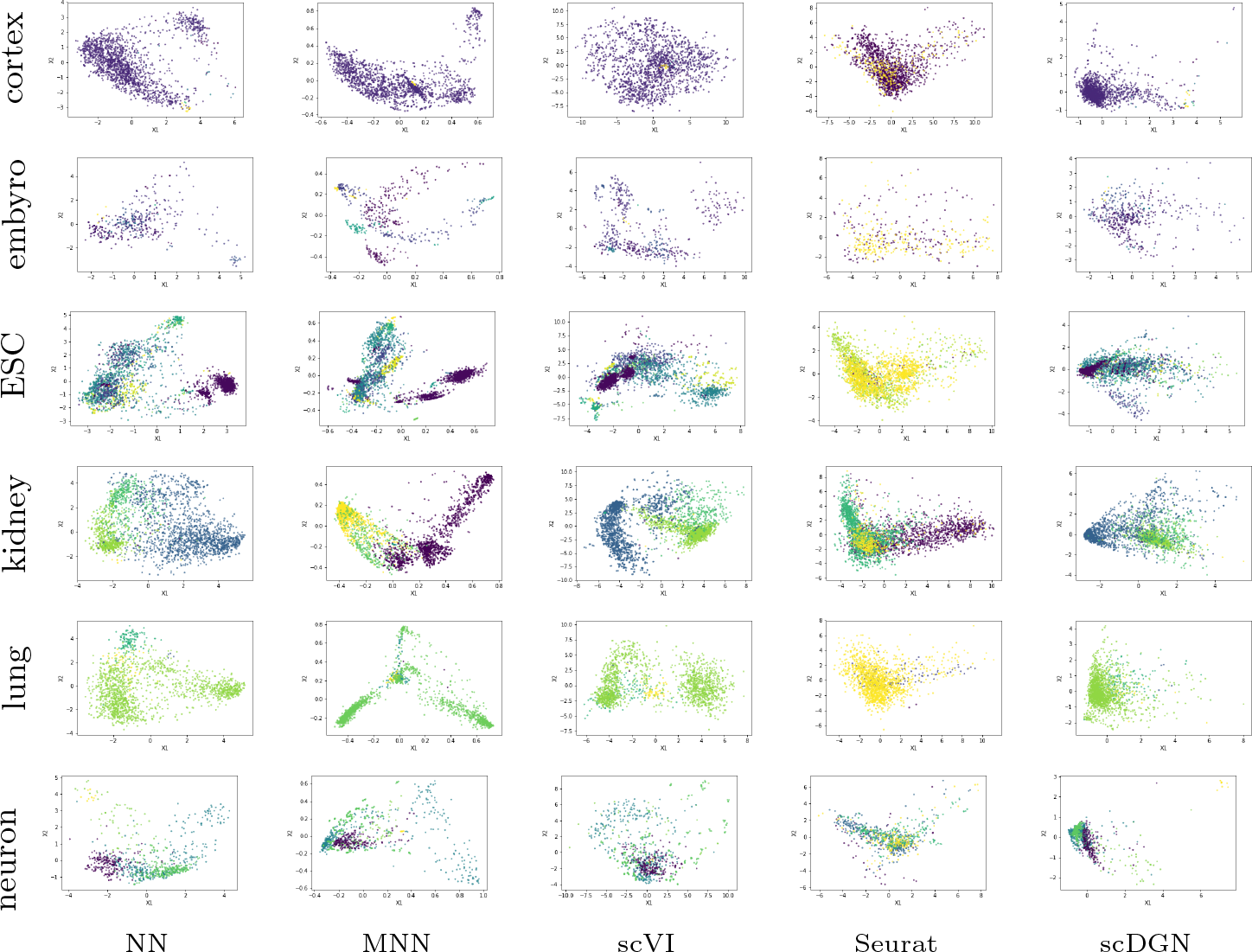
PCA visualization of the representations learned by different models on scQuery dataset for certain cell types. The colors are used to distinguish the batches.

**Fig. C.10:**
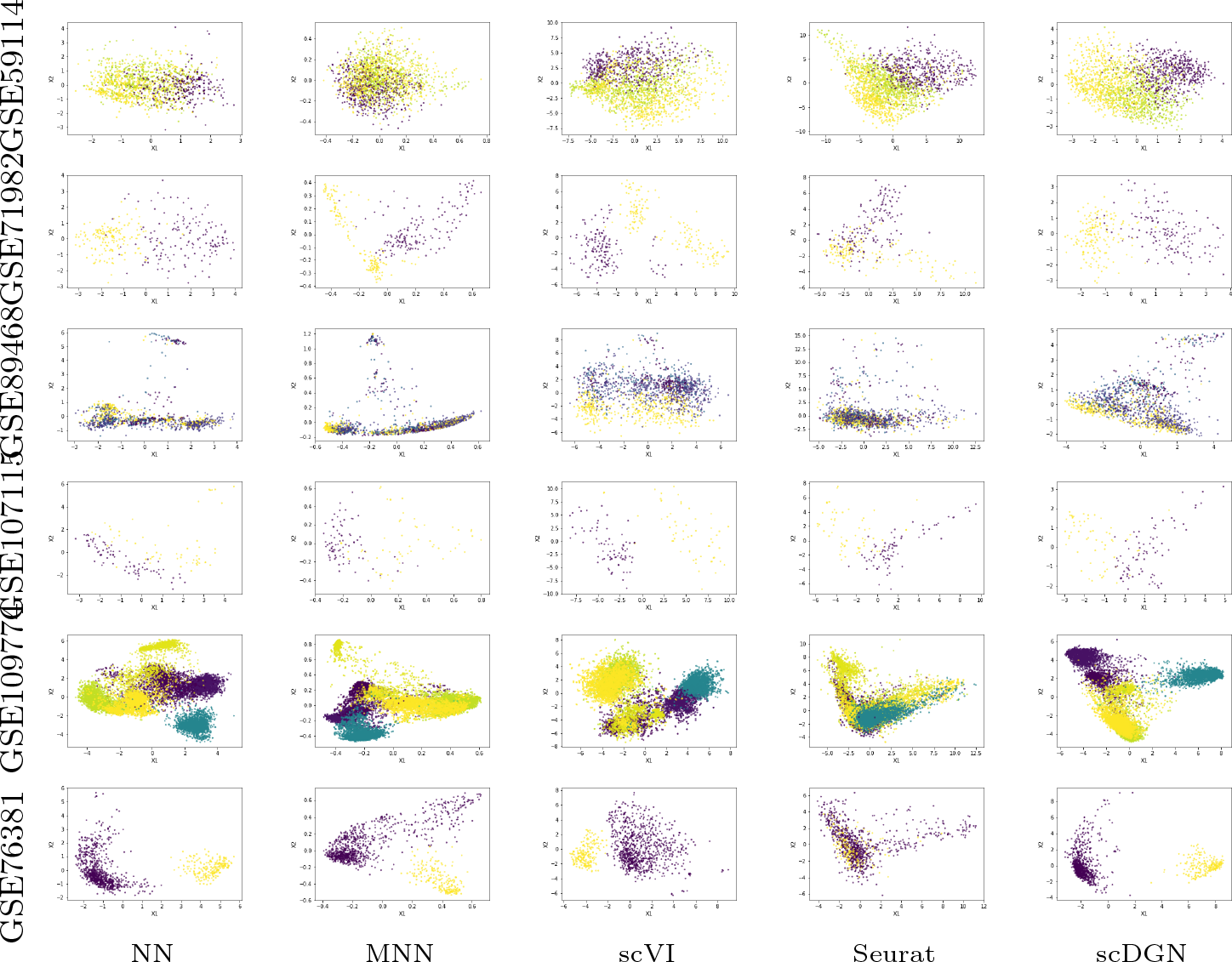
PCA visualization of the representations learned by different models on sc-Query dataset for certain batches. The colors are used to distinguish the cell types.

### C.2 Full Pancreas Datasets

**Fig. C.11:**
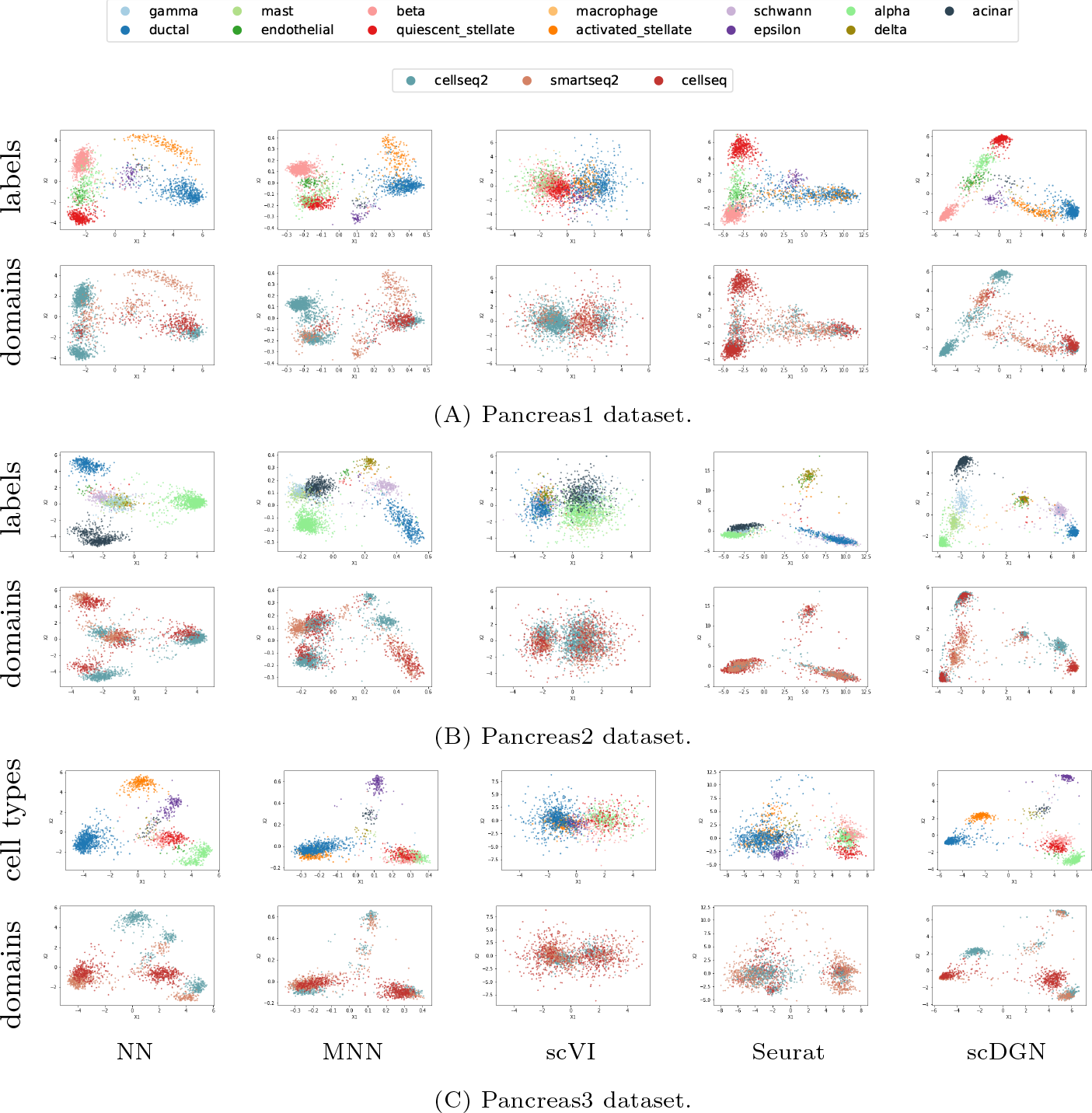
PCA visualization of the representations learned by different models on the whole Pancreas datasets.

**Fig. C.12:**
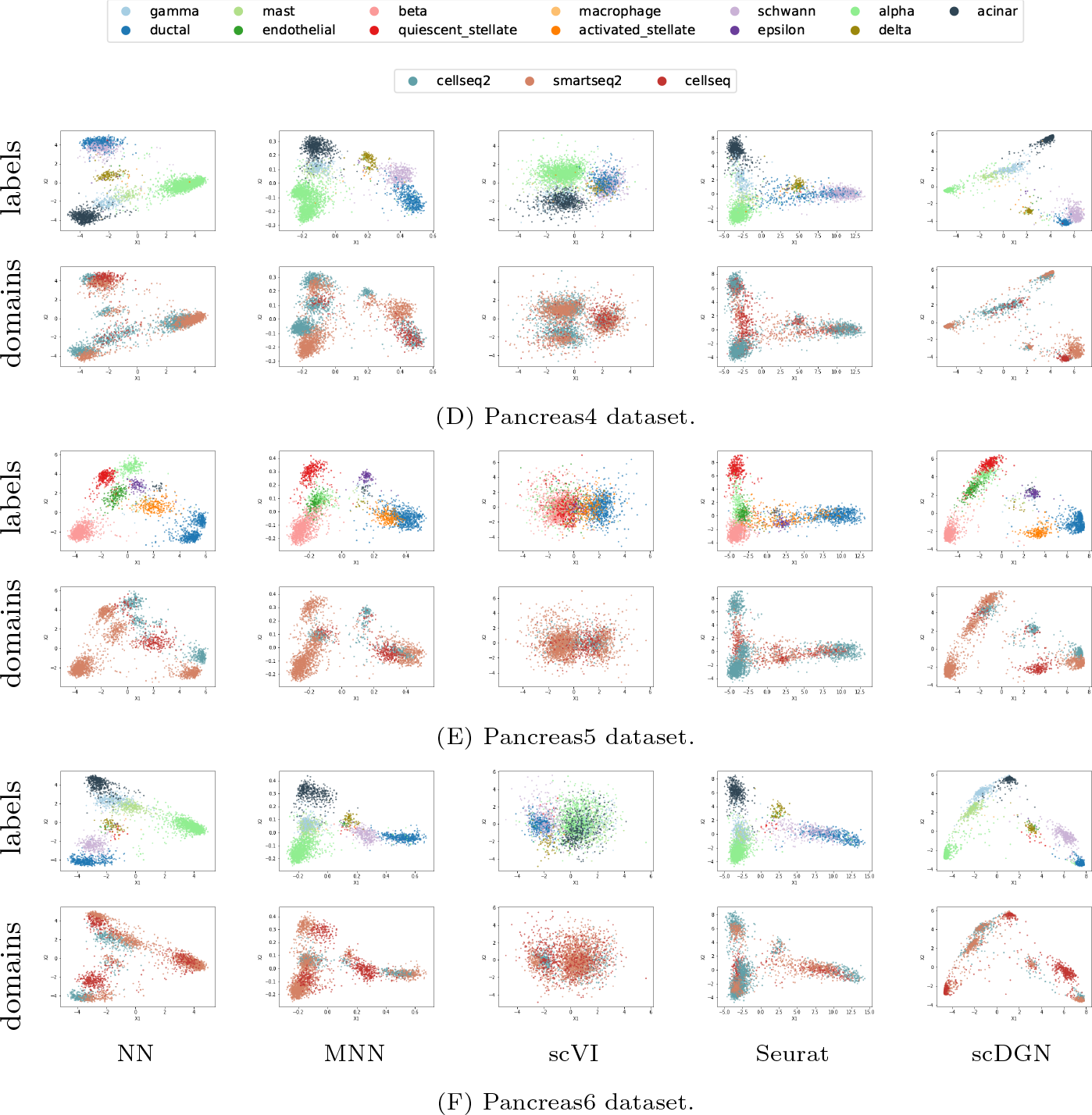
PCA visualization of the representations learned by different models on the Pancreas datasets.

### C.3 Pancreas1

**Fig. C.13:**
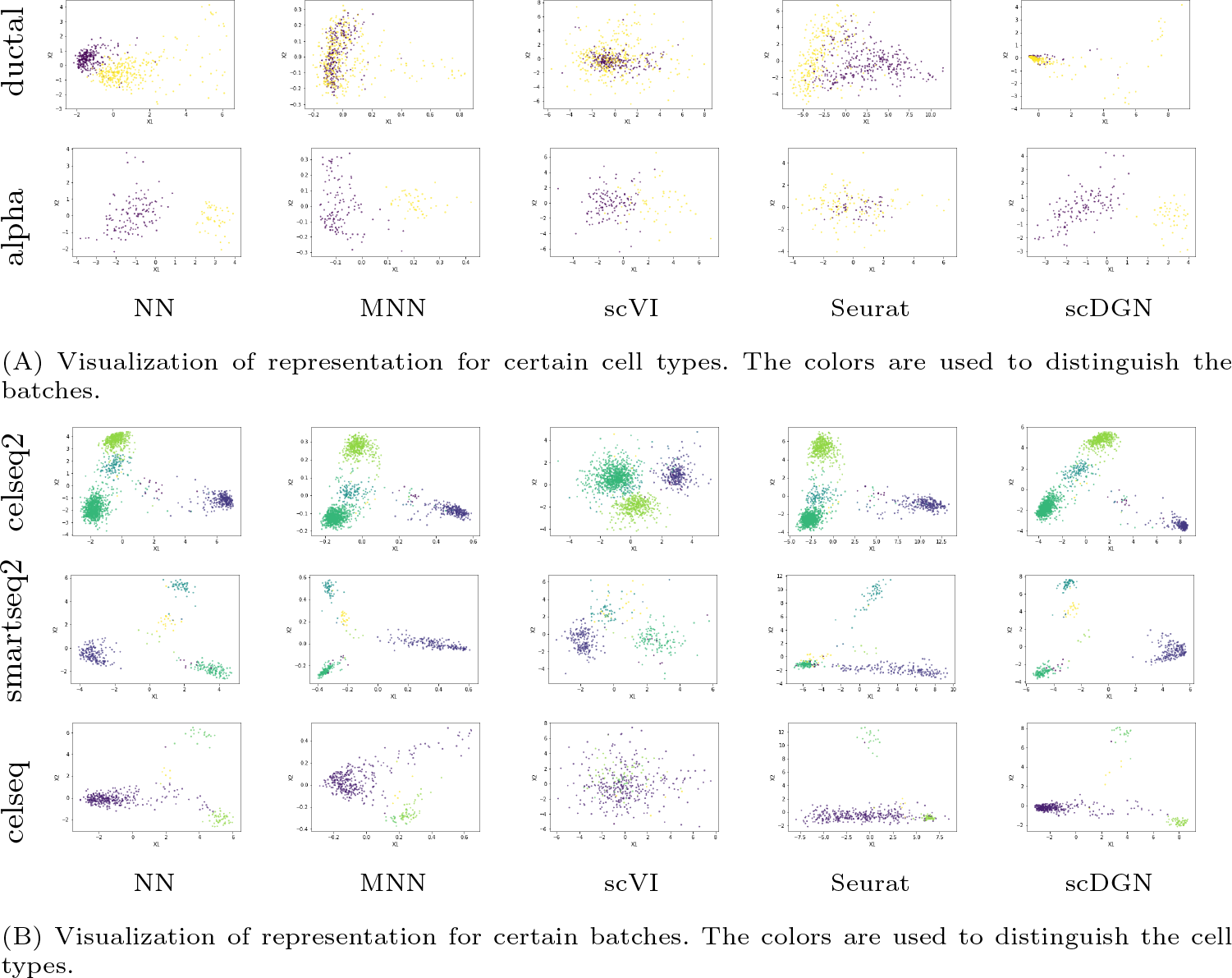
PCA visualization of the representations for certain cell types and domains on Pancreas1 dataset.

### C.4 Pancreas2

**Fig. C.14:**
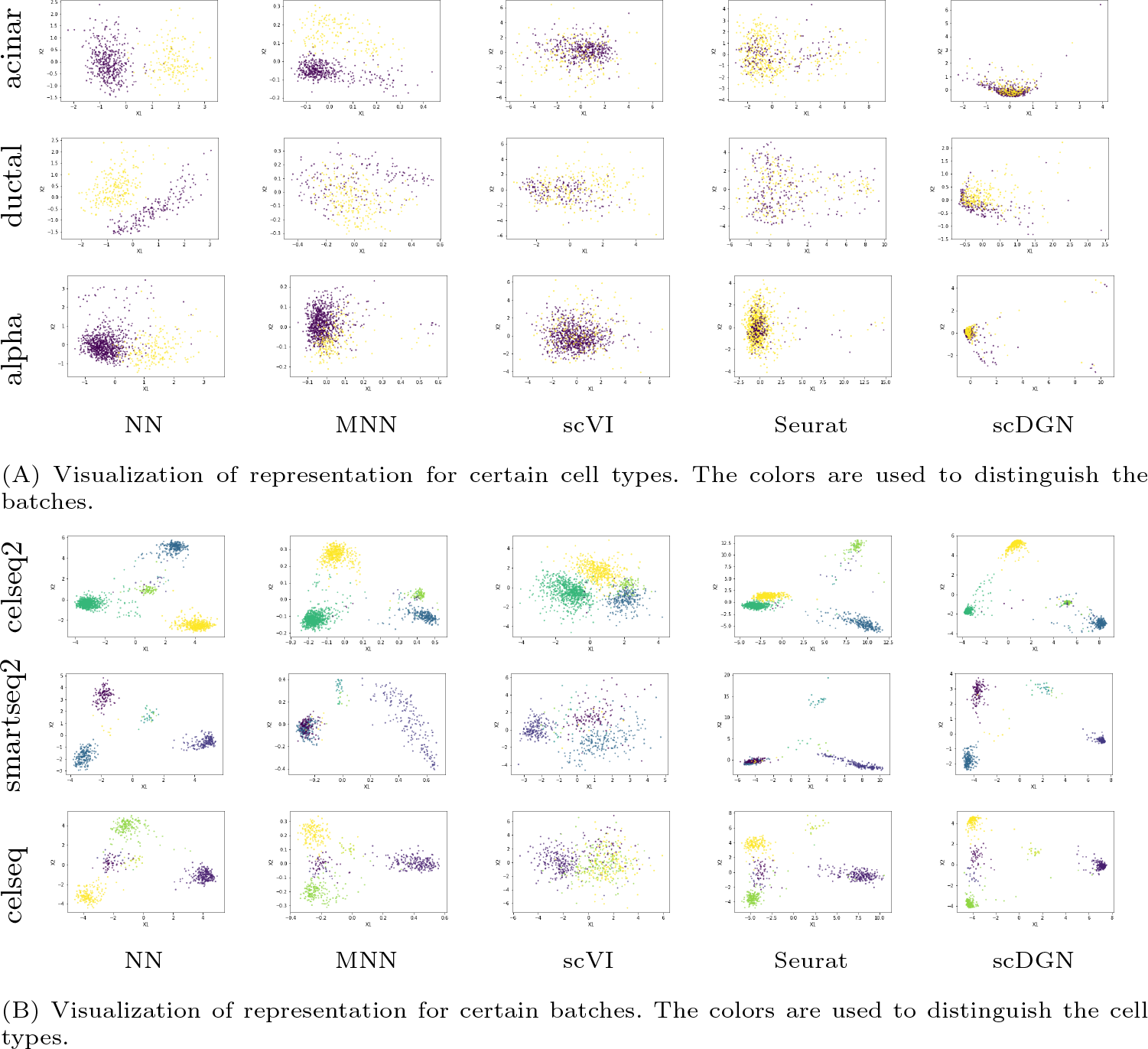
PCA visualization of the representations for certain cell types and domains on Pancreas2 dataset.

### C.5 Pancreas3

**Fig. C.15:**
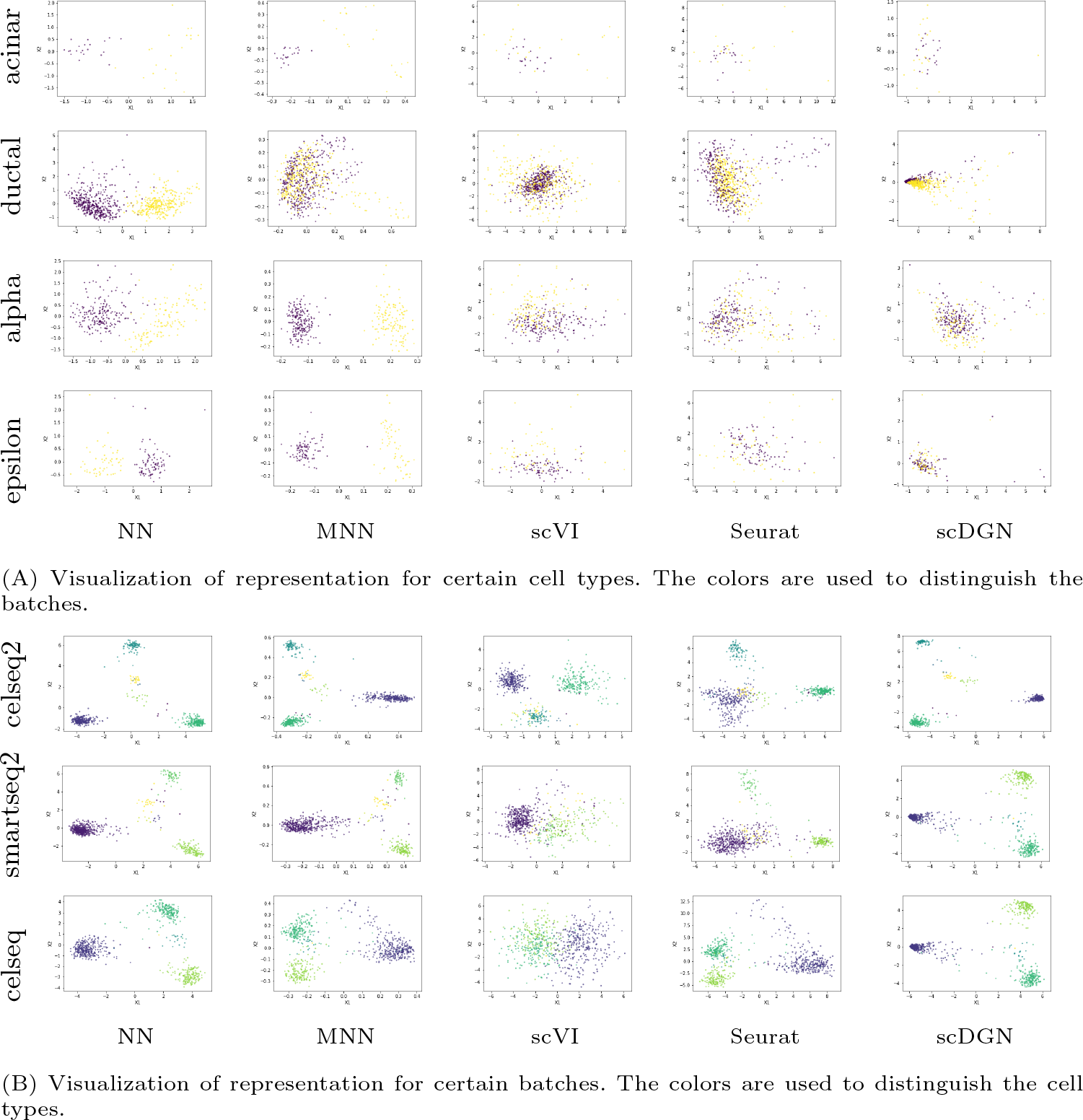
PCA visualization of the representations for certain cell types and domains on Pancreas3 dataset.

### C.6 Pancreas4

**Fig. C.16:**
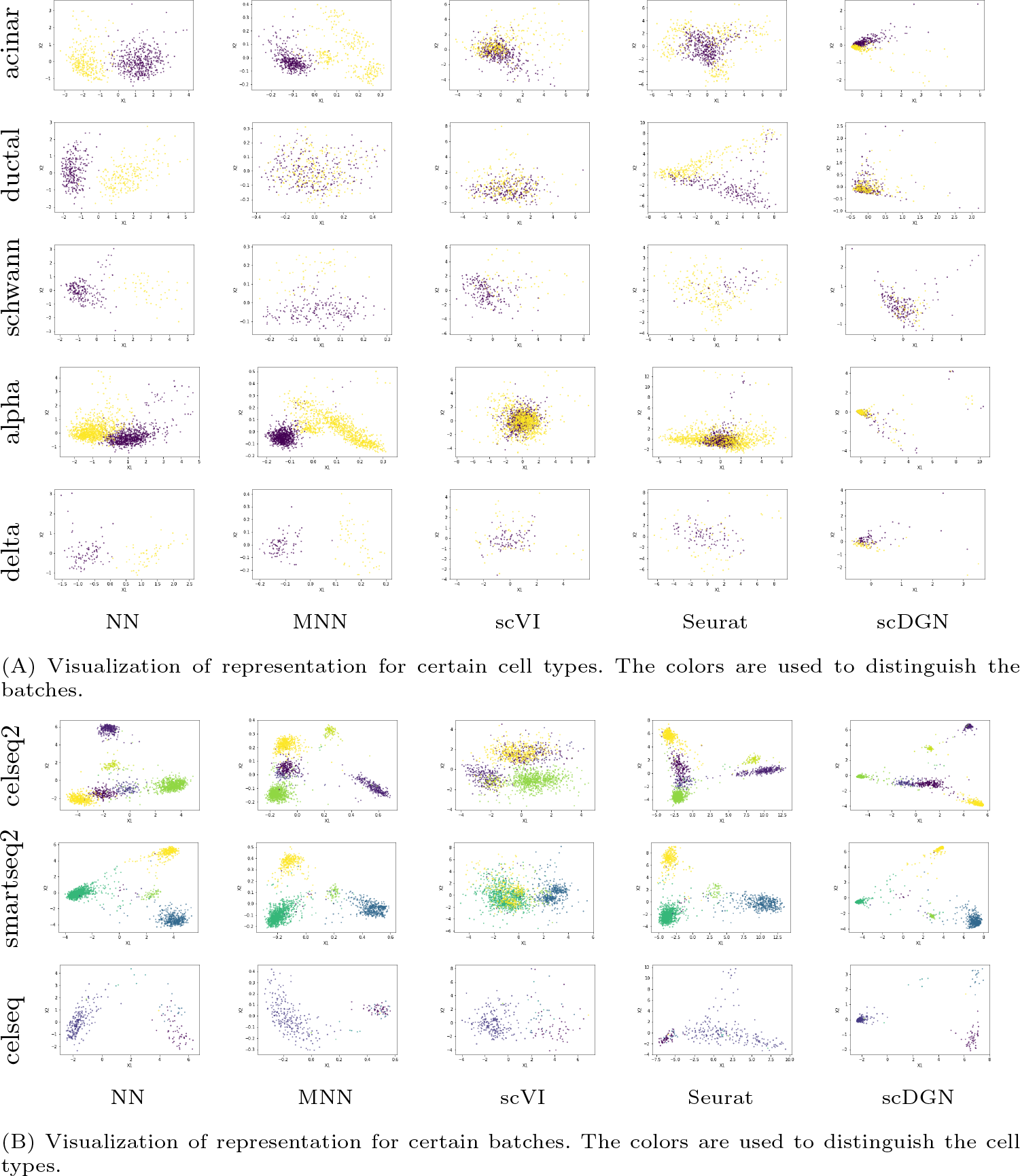
PCA visualization of the representations for certain cell types and domains on Pancreas4 dataset.

### C.7 Pancreas5

**Fig. C.17:**
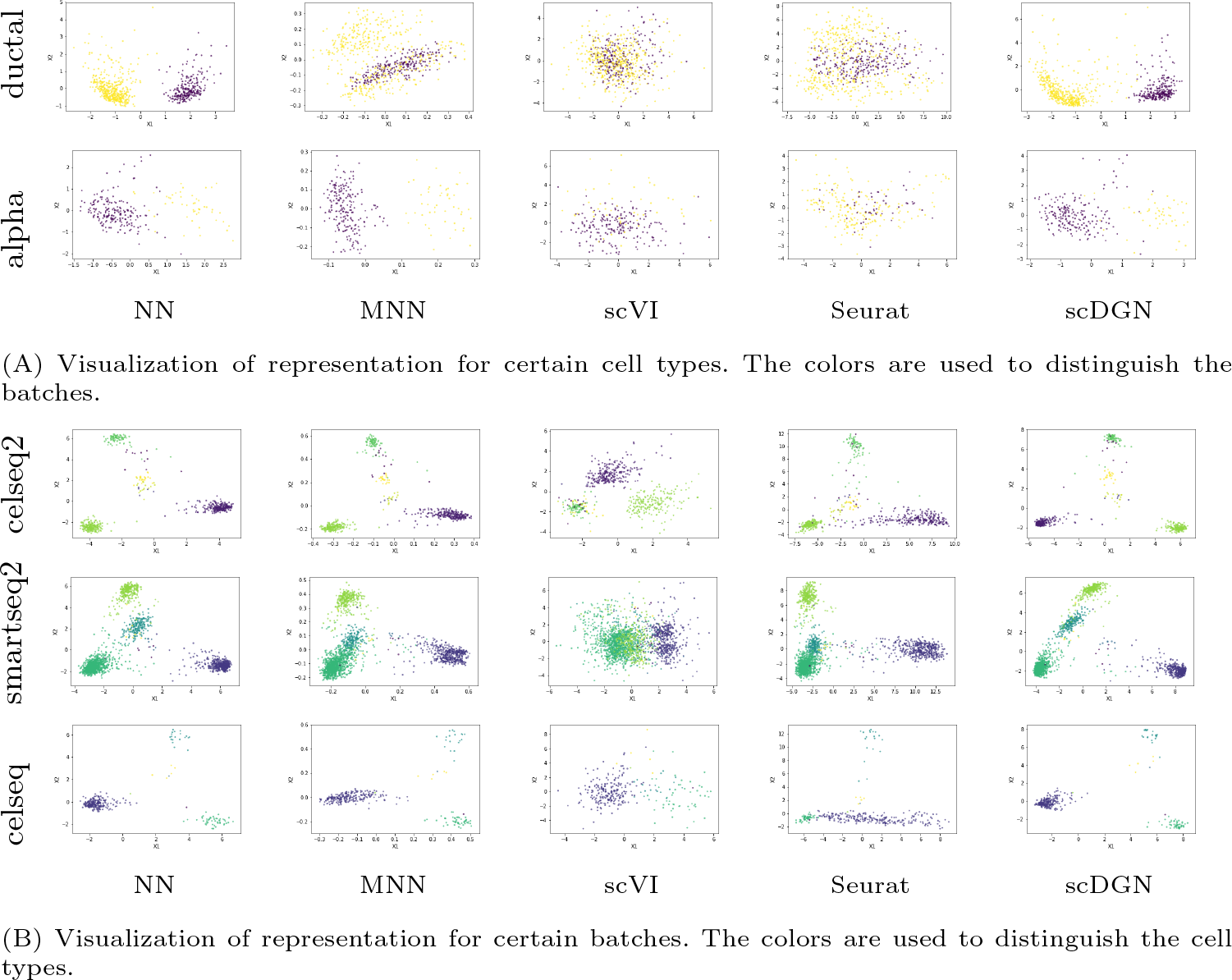
PCA visualization of the representations for certain cell types and domains on Pancreas5 dataset.

### C.8 Pancreas6

**Fig. C.18:**
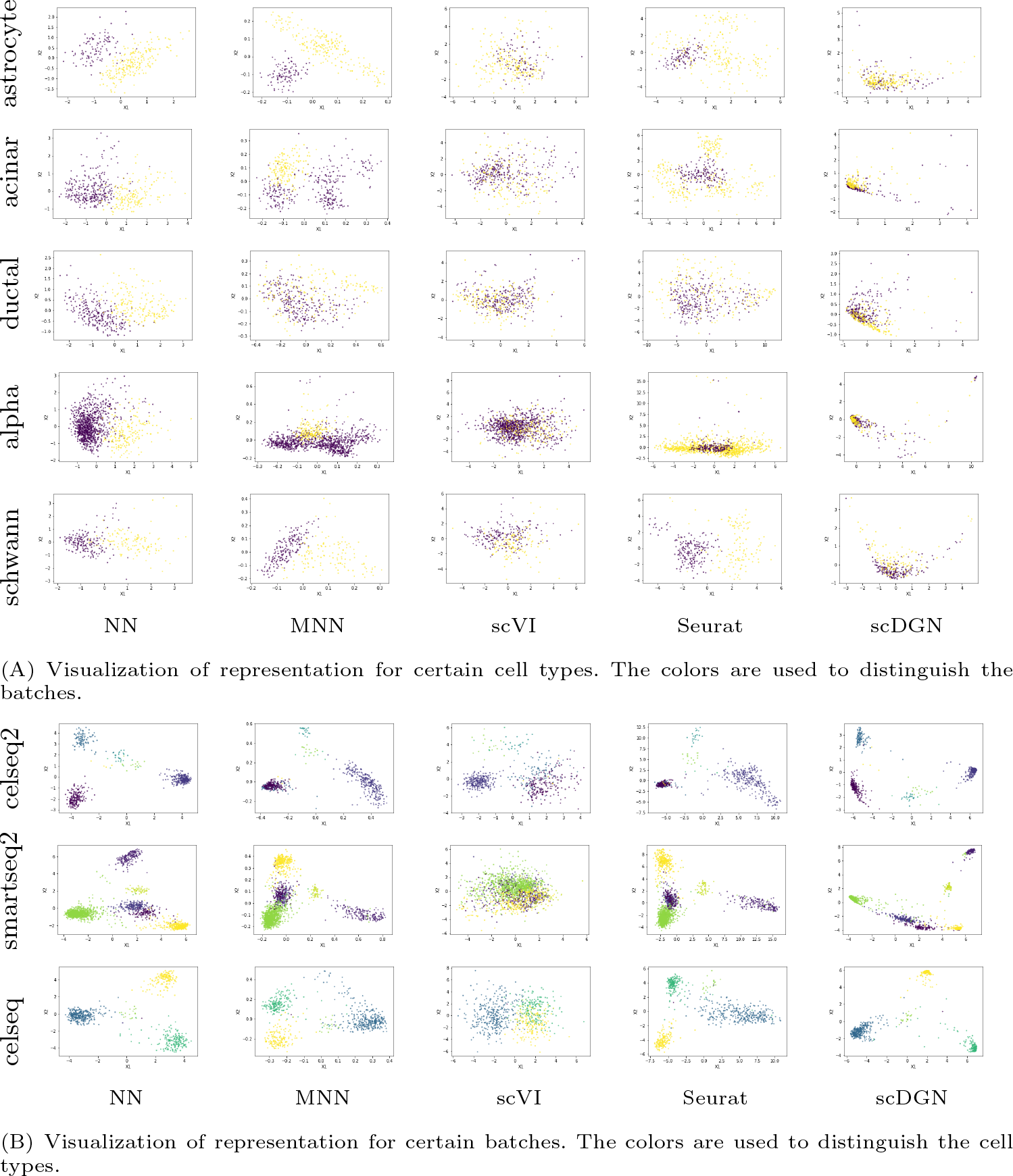
PCA visualization of the representations for certain cell types and domains on Pancreas6 dataset.

## D Key gene analysis

### D.1 Top 100 Genes for Liver Cell Type^2^

**– NN:** ubb, *ins2*, *apoe*, *hba-a1*, *actb*, *b2m*, *hba-a2*, *mup11*, *cd74*, *dnase1l3*, *h2-d1*, *mup22*, *tuba1a*, *tmsb4x*, *mup14*, *selenop*, *aw112010*, *ins1*, *cst3*, *mup7*, *iapp*, *ccdc152*, *ftl1*, *mup13*, *mup10*, *chl1*, *hbb-bt*, *clec4g*, *igfbp7*, *nrep*, *mup19*, *sh3bgrl3*, *hbb-bs*, *mup1*, *mup18*, *h2-ab1*, *mup16*, *mup12*, *rpl18a*, *c1qb*, *map1lc3b*, *mup15*, *ccl5*, *hbb-y*, *rps29*, *ctsd*, *ly86*, *ifitm2*, *mup9*, *h3f3b*, *mup8*, *nkg7*, *tmsb10*, *mt1*, *tagln2*, *ttr*, *gstm1*, *hba-x*, *atpif1*, *lyz2*, *apoc1*, *lgals1*, *mup2*, *stmn2*, *apoa2*, *plp1*, *calm1*, *gpx3*, *prdx1*, *igfbp4*, *prl3d1*, *c1qa*, *apod*, *tma7*, *h2-aa*, *cmss1*, *ptma*, *cd79a*, *gapdh*, *plpp1*, *sst*, *abi1*, *dennd1b*, *cfl1*, *rgs1*, *gclm*, *pitpnc1*, *dppa3*, *fcgr2b*, *serp1*, *rpl8*, *tyrobp*, *sumo1*, *zfp976*, *rplp1*, *gm10591*, *gm21541*, *rpsa*, *s100a8*, *pcnp*.

**– scDGN:** rps29, *apoe*, *mup11*, *ins2*, *mup22*, *dnase1l3*, *selenop*, *mup10*, *b2m*, *gcg*, *mup7*, *h2-d1*, *mup14*, *tmsb10*, *ccdc152*, *clec4g*, *apoa2*, *hba-a1*, *ttr*, *mup13*, *myl7*, *mgp*, *mup18*, *mup16*, *mt1*, *mup15*, *npm1*, *mup19*, *hba-a2*, *aw112010*, *igfbp7*, *chl1*, *mup1*, *actg1*, *apoc1*, *ins1*, *ly6e*, *mup9*, *mup8*, *ptma*, *mup12*, *mup2*, *cst3*, *fabp3*, *btg1*, *iapp*, *rpl35*, *apoa1*, *h2-k1*, *cmss1*, *mup17*, *lgals1*, *tmsb4x*, *stmn1*, *gm13304*, *ptp4a2*, *prdx1*, *gm21541*, *hmga1*, *snap25*, *set*, *plp1*, *ccl4*, *fabp4*, *trim30a*, *gstp1*, *gm10591*, *ubc*, *scgb1a1*, *resp18*, *sumo1*, *fabp1*, *nrep*, *h2-q7*, *npy*, *itm2b*, *hspe1*, *car2*, *sub1*, *slc25a5*, *h2afz*, *ywhah*, *ccl21b*, *pomc*, *rpl41*, *cbx3*, *ctsd*, *rps27rt*, *laptm5*, *chchd2*, *s100a8*, *actc1*, *hba-x*, *hbb-bt*, *myl4*, *eif1*, *gpihbp1*, *sod1*, *gabarapl2*, *calm1*.

### D.2 GO Analysis Results for the Other Cell Types

**Table 5:** GO analysis results of the genes with respect to embryonic stem cell that are only recognized by scDGN.

| term_name | term_id | $p_{adj}$ | $-\log_{10} p_{adj}$ |
| --- | --- | --- | --- |
| SRP-dependent cotranslational protein targeting to membrane | GO:0006614 | 1.8109E-06 | 5.74210612 |
| cotranslational protein targeting to membrane | GO:0006613 | 2.59394E-06 | 5.58603936 |
| protein targeting to ER | GO:0045047 | 4.44101E-06 | 5.35251780 |
| establishment of protein localization to endoplasmic reticulum | GO:0072599 | 5.72221E-06 | 5.24243629 |
| nuclear-transcribed mRNA catabolic process, nonsense-mediated decay | GO:0000184 | 9.79366E-06 | 5.00905515 |
| protein localization to endoplasmic reticulum | GO:0070972 | 2.19947E-05 | 4.65768238 |

**Table 6:** GO analysis results of the genes with respect to hematopoietic stem cell that are only recognized by scDGN.

| term_name | term_id | $p_{adj}$ | $-\log_{10} p_{adj}$ |
| --- | --- | --- | --- |
| protein targeting to ER | GO:0045047 | 0.041444862 | 1.382529302 |
| cotranslational protein targeting to membrane | GO:0006613 | 0.026817903 | 1.571575182 |
| SRP-dependent cotranslational protein targeting to membrane | GO:0006614 | 0.020030591 | 1.698306246 |
| protein localization to endoplasmic reticulum | GO:0070972 | 0.013637519 | 1.865264616 |
| translational initiation | GO:0006413 | 0.001555052 | 2.808254977 |

## Footnotes

1 https://scquery.cs.cmu.edu/processed_data/

2 The results in text format are also available at: https://github.com/SongweiGe/scDGN/blob/master/supplementarymaterials/top_genes.txt

